# INNATE IMMUNE INTERFERENCE ATTENUATES INFLAMMATION IN *BACILLUS* ENDOPHTHALMITIS

**DOI:** 10.1101/2020.06.10.144915

**Authors:** Md Huzzatul Mursalin, Phillip S. Coburn, Frederick C. Miller, Erin Livingston, Roger Astley, Michelle C. Callegan

## Abstract

**PURPOSE:** *Bacillus* endophthalmitis is a sight-threatening bacterial infection that sometimes requires enucleation. Inflammation in this disease is driven by activation of innate Toll-like receptor (TLR) pathways. Here, we explored the consequences of innate immune interference on intraocular inflammatory responses during *Bacillus* endophthalmitis.

**METHODS:** Endophthalmitis was induced in mice by injecting 100 CFU *Bacillus thuringiensis* in to the mid-vitreous. We interfered with activation of the TLR2 and TLR4 pathways by 1) injecting a group of mice with S layer protein-deficient (Δ*slpA*) *B. thuringiensis* or 2) injecting a group of wild type (WT)-infected mice with a TLR2/4 inhibitor, oxidized phospholipid (OxPAPC). At 10 hours postinfection, infected eyes were removed and total RNA was purified. mRNA expression was then analyzed by NanoString using a murine inflammation panel. We compared findings with expression data from eyes infected with eyes injected with WT *B. thuringiensis*, eyes injected with OxPAPC alone, and uninfected eyes.

**RESULTS:** Interference of TLR2 and TLR4 pathways resulted in differential expression of mouse inflammatory genes compared to expression in WT-infected eyes. In WT-infected eyes, 56% of genes were significantly upregulated compared to that of uninfected controls. However, compared to WT-infected eyes, the expression of 27% and 50% of genes were significantly reduced in WT+OxPAPC and Δ*slpA-*infected eyes, respectively. The expression of 61 genes which were significantly upregulated in WT-infected eyes was decreased in WT+OxPAPC or Δ*slpA-*infected eyes. Interference with activation of the TLR2 and TLR4 pathways resulted in blunted expression of complement factors (C3, Cfb, and C6) and several innate genes such as TLR2, TLR4, TLR6, TLR8, MyD88, Nod2, Nlrp3, NF-κB, STAT3, RelA, RelB, and Ptgs2. Interference with activation of the TLR2 and TLR4 pathways also reduced the expression of several inflammatory cytokines such as CSF3, IL-6, IL-1β, CSF2, IL-1α, TNFα, IL-23α, TGFβ1, and IL-12β and chemokines CCL2, CCl3, CXCL1, CXCL2, CXCL3, CXCL5, CXCL9, and CXCL10. All of the aforementioned genes were significantly upregulated in WT-infected eyes.

**CONCLUSIONS:** These results suggest that interfering with the activation of innate immune pathways during *Bacillus* endophthalmitis significantly reduced the intraocular inflammatory response. This positive clinical outcome could be a strategy for anti-inflammatory therapy of an infection typically refractory to corticosteroid treatment.

## INTRODUCTION

Bacterial endophthalmitis is a dangerous ocular infection that is considered a medical emergency. This intraocular infection is often caused by Gram-positive bacteria ^1–4^. The infecting organism can be introduced into the immune-privileged posterior chamber of the eye by an injury (post-traumatic), after a surgical procedure (post-operative), or by metastasis of bacteria into the eye from a distant infection site (endogenous) ^5–8^. Regardless of the route of infection, the signs and symptoms are quite similar, ranging from red eyes and swollen eyelids to severe intraocular pain and vision loss ^1–4^. The severity of the disease can range from mild inflammation that resolves with treatment, to a fulminant, rapidly progressing infection that is refractory to currently available treatment options. *Bacillus* causes the most severe form of bacterial endophthalmitis and is often associated with worse patient outcomes than infections caused by other pathogenic species ^6,8,9–11^. The pathogenicity of *Bacillus* endophthalmitis is associated with the inflammogenicity of the bacterial cell wall and the production of secreted toxins and enzymes ^12–18^. The very severe nature of *Bacillus* endophthalmitis dictates the need for prompt and rapid therapy to stop the disease progression. However, *Bacillus* endophthalmitis may result in vision loss within 12-24 hours of infection, despite therapeutic intervention that may otherwise effectively attenuate infection caused by other ocular pathogens such as *Staphylococcus aureus* or *Streptococcus pneumoniae* ^9^.

*Bacillus* is a Gram-positive, motile, spore-forming rod, and is distributed throughout nature, especially in soil ^19–21^. Metabolically-inactive *B. cereus* triggered an inflammatory response in a rabbit experimental endophthalmitis model, suggesting a significant role of cell wall components in inciting inflammation ^11^. In addition to the thick outer layer containing peptidoglycan, lipoteichoic acid, and lipoproteins, the unique architecture of the *Bacillus* envelope also includes flagella and a monomolecular crystalline array of proteinaceous subunits termed surface layers or S-layer ^22–25^. If present, S-layer protein (SLP) is one of the most abundant proteins in the bacterial cell envelope and provides the organism with a selective advantage in diverse habitats ^26^. SLP contributes to colonization by promoting bacterial adherence and biofilm formation ^27,28^, as well as protecting the bacteria against complement killing and phagocytosis ^29–32^. We recently demonstrated that the absence of SlpA resulted in significant retention of retinal function, reduced inflammatory cell influx, and muted disease severity in a mouse experimental endophthalmitis model of *Bacillus* endophthalmitis ^33^. We also reported *Bacillus* SlpA as a stimulator of nuclear factor kappa-light-chain-enhancer of activated B cells (NF-κB) in human retinal Muller cells *in vitro* ^34^. SLP also protects *Bacillus* from phagocytosis by neutrophils and retinal cells and also impacts bacterial adherence to retinal cells ^34^. SLP appears to be an important virulence factor in the pathogenesis of *Bacillus* endophthalmitis.

During infection, ocular immune privilege ^35^ is compromised by an overwhelming, acute inflammation resulting from interactions between innate receptors and microbial ligands. We reported that Toll-like receptors (TLR) 2 and 4, but not TLR5, were essential for rapid intraocular inflammation during *Bacillus* infection ^36–38^. Being a Gram-positive pathogen, *Bacillus* does not produce lipopolysaccharide (LPS), so the specific ligand on *Bacillus* that interacts with TLR4 remained unknown. We reported that *Bacillus* SLP activated both TLR2 and TLR4 ^34^. When activation of these TLRs was blocked by injection of oxidized phospholipid (OxPAPC) during experimental endophthalmitis, disease severity and overall inflammation were reduced and retinal function was retained compared to that in untreated control mice ^34^. OxPAPC is produced by the oxidation of 1-palmitoyl-2-arachidonyl-sn-glycero-3-phosphorylcholine (PAPC) and has been shown to inhibit the signaling induced by bacterial lipopeptide and lipopolysaccharide (LPS), known agonists for TLR2 and TLR4. OxPAPC competes with CD14, lipid binding protein (LBP), and MD2, and thus blocks signaling in TLR2 and TLR4 pathways ^39,40^.

TLR-ligand interactions during intraocular infection generates inflammatory mediators that recruit neutrophils into the eye ^41,42^. When TLRs recognize surface molecules on pathogenic bacteria, signals are transmitted via the adaptor molecules to the signaling molecules which are located in the cytoplasm. Signals from cytoplasm reach the nucleus and activate the transcription of inflammatory mediators. We reported that both myeloid differentiation primary response gene-88 (MyD88) and Toll/interleukin-1 receptor (TIR) domain-containing adaptor-inducing interferon-β (TRIF) regulate this inflammation signaling cascade during murine experimental endophthalmitis ^38^. Neutrophils are the primary innate responders and may cause bystander damage to the retina during an intraocular infection ^43–45^. Therefore, inflammatory mediators that recruit neutrophils are crucial for disease outcome. During experimental *Bacillus* endophthalmitis, the absence of tumor necrosis factor (TNF) α resulted in reduced neutrophil infiltration into the eye, which resulted in an elevated bacterial load and damage to ocular tissues ^44^. The absence of TNFα was compensated for by elevated expression of other mediators such as chemokine (C-X-C motif) ligand (CXCL) 1, CXCL2/MIP1α, and IL-6, which might have contributed to delayed recruitment of neutrophils and overall pathogenesis ^44^. We reported that the absence of CXCL1 or anti-CXCL1 treatment led to a blunted inflammatory response but improved clinical outcome, suggesting a potential benefit for targeting neutrophil chemoattractants to curb inflammation-mediated damage ^45^.

In endophthalmitis, the use of anti-inflammatory therapeutics with antibiotics has not proven completely effective at improving disease outcomes ^5–8^. The primary function of the innate inflammatory response is to detect invading pathogens and clear them as quickly as possible. However, the highly inflammogenic *Bacillus*-induced ocular inflammatory response is often so robust that it is difficult to control and ultimately results in vision loss ^9,46^. At present, a universal therapeutic regimen to prevent vision loss in *Bacillus* endophthalmitis is not available. Since the depletion of individual proinflammatory mediators could be compensated for by the synthesis of other mediators, finding anti-inflammatory agents with the potential to block multiple inflammatory pathways might hold the potential for future anti-inflammatory therapeutics.

An acute and potentially damaging inflammatory response in the eye is one of the major problems of bacterial endophthalmitis. We reported that an absence of TLR2 or TLR4 in knockout mice or inhibition of TLR2 and TLR4 activation by OxPAPC improved the clinical outcome of experimental *Bacillus* endophthalmitis ^33,36,37^. OxPAPC-mediated interference of innate activation in response to pathogens and pathogen associated molecular patterns (PAMPs) are well recognized ^39,40,47,48^. In this study, we investigated how interfering with innate immune pathways during the early stage of *Bacillus* infection would impact inflammation during the later stage of infection. Since inhibition of TLR signaling serves as a promising treatment option for inflammatory diseases ^49^, we hypothesized that innate interference could be protective for a rapidly blinding disease like *Bacillus* endophthalmitis. Innate pathway interference was achieved by infection with SLP-deficient *Bacillus* (Δ*slpA*) or treatment of experimental *Bacillus* endophthalmitis with OxPAPC. NanoString analysis of a mouse inflammation panel revealed that innate interference considerably reduced inflammatory gene expression in the whole infected eye. Our findings suggest that the attenuation of innate activation may be a viable anti-inflammatory option for the treatment of *Bacillus* and perhaps other types of endophthalmitis.

## MATERIALS AND METHODS

### Ethical statement

The experiments in this study involved the use of mice. All animal experiments were performed in strict accordance with the recommendations in the Guide for the Care and Use of Laboratory Animals and the Association for Research in Vision and Ophthalmology Statement for the Use of Animals in Ophthalmic and Vision Research. The protocols were approved by the Institutional Animal Care and Use Committee of the University of Oklahoma Health Sciences Center (protocol numbers 15-103, 18-043, and 18-087).

### Bacterial strains

*B. thuringiensis* subsp. Galleriae NRRL 4045 (WT) or its isogenic S-layer protein-deficient mutant (Δ*slpA*) was used to initiate experimental endophthalmitis in mice, as previously described ^33,34^. WT *B. thuringiensis* and Δ*slpA B. thuringiensis* were grown to early stationary phase in brain heart infusion (BHI; VWR, Radnor PA) broth for 18 hours and diluted to 100 CFU/0.5 μl for injection into the mid-vitreous.

### Mice and Intraocular Infection

C57BL/6J mice purchased from Jackson Laboratories (Bar Harbor ME, Stock No. 000664) were used for all experiments. Upon arrival, they were housed on a 12 hour on/12 hour off light cycle in biohazard level 2 conditions and acclimated for at least two weeks to equilibrate their microbiota. Mice were 8-10 weeks of age at the time of the experiments. A combination of ketamine (85 mg/kg body weight; Ketasthesia, Henry Schein Animal Health, Dublin, OH) and xylazine (14 mg/kg body weight; AnaSed, Akorn Inc., Decatur, IL) were used to sedate the mice. Four groups of C57BL/6J mice with three mice in each group were used in these experiments (Figure 1A). The first two groups were infected with 100 CFU WT *B. thuringiensis*/0.5 μl BHI, and the third group was infected with 100 CFU Δ*slpA B. thuringiensis*/0.5 μl BHI,as previously described ^33,34^. At 4 hours postinfection, the second group of WT *B. thuringiensis*-infected mice was intravitreally treated with the synthetic TLR2/4 inhibitor OxPAPC (Invivogen; 30 ng/μl) (WT+OxPAPC) ^34^. Uninfected mice were used as control.

**Figure 1.**
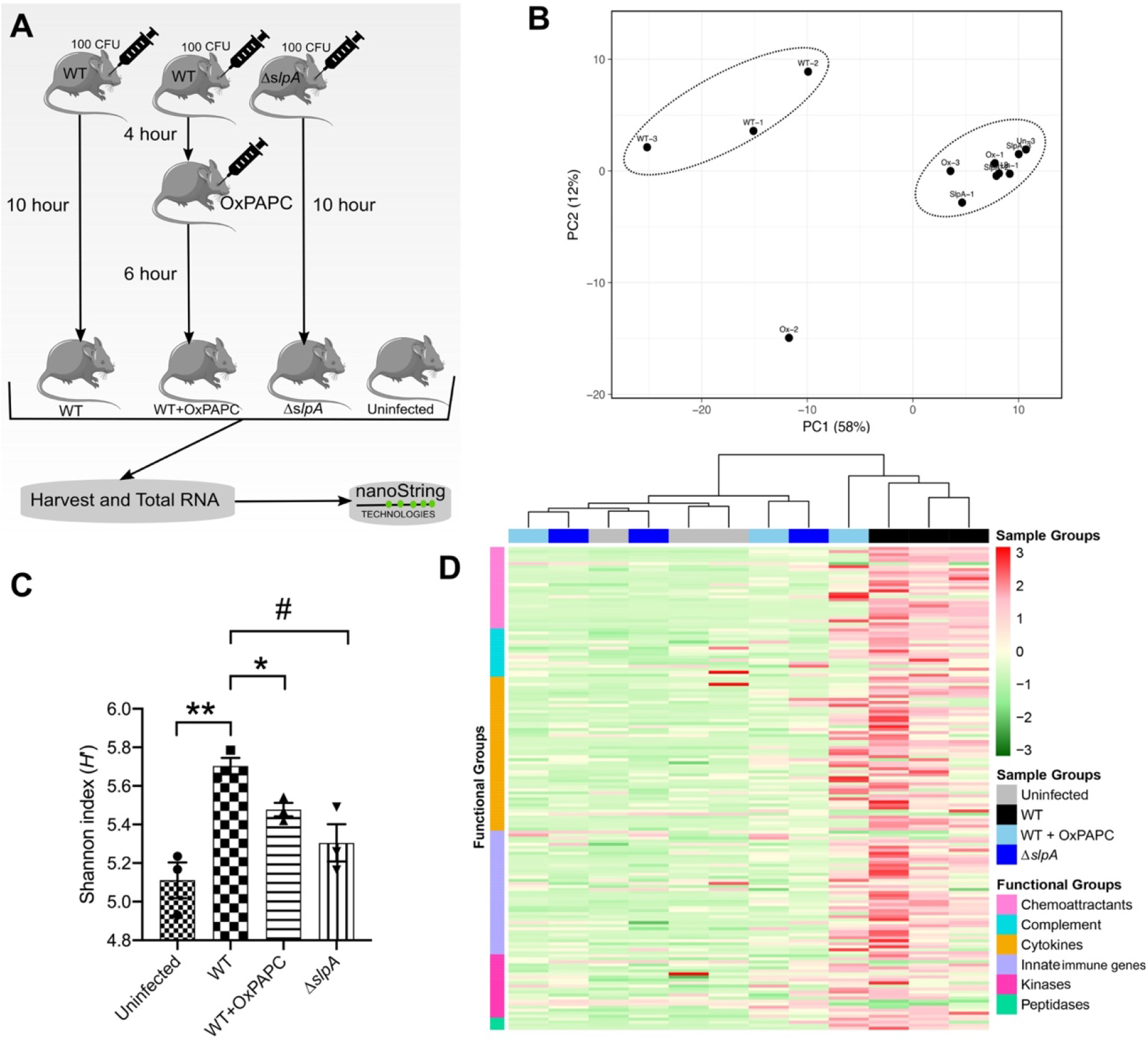
Interfering with TLR2 and TLR4 activation affected inflammatory gene expression in the eye. (A) Experimental outline of the NanoString experiment. (B) PCA analysis of mouse inflammatory gene. All WT-infected eyes cluster together. Except for one WT+OxPAPC eye, all uninfected, Δ*slpA*-infected, and two WT+OxPAPC eyes clustered together. (C) Variations in the diversity of gene expression profiles across different groups. Shannon index was used to calculate the diversity. Statistical significance was calculated using unpaired t-test. *P=0.0152, **P=0.0043, and #P=0.0198. (D) Heatmap for the visualization of gene expression. The red and green color gradient represents the upregulation and downregulation of genes. Red indicates increased, whereas green represents reduced gene expression. Sample groups were color-coded by grey (uninfected), black (WT-infected), light blue (WT+OxPAPC), and deep blue (Δ*slpA*-infected). Functional groups were color-coded in the Y axis as follows: chemoattractants (light pink), complement (light blue), cytokines (yellow), innate elements (purple), kinases (deep pink), and peptidases (green).

### Harvesting mouse eyes and RNA preparation

At 10 hours postinfection, all mice were euthanized by CO2 inhalation. Experimental and control eyes were harvested and transferred to individual 1.5ml screw-cap tubes containing sterile 1mm glass beads (BioSpec Products, Inc., Bartlesville OK) and 400μl lysis buffer (RLT) from an RNeasy kit (Qiagen, Germantown, MD). Eyes were homogenized as previously described using a Mini-BeadBeater (Biospec Products Inc., Bartlesville, OK) for 120 seconds (2 pulses of 60 seconds each). Eye homogenates were centrifuged and transferred into another screw-cap tube containing 0.1 mm glass beads (Biospec Products Inc., Bartlesville, OK) and homogenized for 60 seconds in the Mini-BeadBeater. Tissue lysates were recovered by centrifugation and processed for total RNA purification using the RNeasy kit according to the manufacturer’s instructions. Genomic DNA was removed using the TURBO DNA-free kit (ThermoFisher Scientific, Inc., Waltham, Massachusetts). Eluted RNA was cleaned and concentrated using RNA Clean & Concentrator (Zymo Research, CA, USA). The concentration and purity were confirmed on a Nanodrop spectrophotometer ^50^.

### NanoString Analysis

In this technique, a biotin-labeled capture probe and a fluorescent-labeled reporter probe hybridize to specific target transcripts. The nCounter NanoString analysis system counts the immobilized RNAs using their barcodes. The NanoString assay was performed using total RNA samples from whole mouse eyes. From each sample, 100ng of RNA was hybridized with NanoString’s XT PGX Mmv2 Inflammation code set containing 248 mouse inflammatory genes and six housekeeping genes using NanoString’s nCounter XT CodeSet Gene Expression Assay Protocol. After hybridization for 16 hours at 65°C, the samples were loaded onto an nCounter Cartridge and run on the NanoString Sprint platform. Data was normalized using NanoString’s nSolver Analysis Software v 4.0. Normalized data of 248 inflammatory genes was then used to analyze the expression profile in WT-infected, WT-infected and OxPAPC-treated (WT+OxPAPC), and Δ*slpA*-infected eyes compared to uninfected eyes.

### Statistics

NanoString analysis was performed in NanoString’s nSolver Analysis Software v 4.0. After data normalization, GraphPad Prism 7 (Graph-Pad Software, Inc., La Jolla, CA) was used for the comparative analysis. Multiple t-tests were performed to compare the means of individual genes from two different groups. Gene expression in uninfected eyes was compared with that of WT-infected, WT+OxPAPC, and Δ*slpA-*infected eyes. We also compared WT-infected eyes with WT+OxPAPC and Δ*slpA*-infected eyes. P values less than 0.05 were considered to be significant.

## RESULTS

### Interfering with TLR2 and TLR4 activation affected inflammatory gene expression in the eye

*Bacillus* induces a robust intraocular inflammatory response that irreversibly damages nonregenerative retinal tissues and results in a devastating clinical outcome. *Bacillus* SLP likely triggers the host inflammatory pathway by activating both TLR2 and TLR4 ^33,34^. To explore the interference of innate activation on intraocular inflammatory responses during the later stages of *Bacillus* endophthalmitis, we performed NanoString analysis on total RNA isolated from our experimental groups. Figure 1A represents the schematics of our experimental approach. In Figure 1B, we performed Principal Component Analysis (PCA) to understand the variability of gene expression in our experimental groups. The normalized mouse inflammatory genes were plotted in ClustVis, a web tool for visualizing the clustering of multivariate data. All replicates of WT-infected eyes clustered together. Except for one eye in the WT+OxPAPC group, replicates of three uninfected, three Δ*slpA*-infected, and two WT+OxPAPC eyes were clustered together, indicating transcriptional similarities among these groups. In Figure 1C, we calculated the Shannon diversity index to understand the diversities of expressed genes between groups. Compared to uninfected eyes, gene expression was more diverse in WT-infected eyes. However, compared to WT-infected eyes, gene expression was less diverse in WT+OxPAPC or Δ*slpA*-infected eyes. In Figure 1D, we employed the R package “pHeatmap” for visualization of gene expression in our groups ^51^. Most of the inflammatory genes in three uninfected, three Δ*slpA*-infected, and two WT+OxPAPC eyes had low levels of expression. In contrast, most of the genes in three WT-infected eyes and one WT+OxPAPC eye had high levels of expression. The branch lengths on the top of the heat map indicate the correlation of gene expression. The longer branches indicate a lower correlation. All uninfected, two WT+OxPAPC, and all Δ*slpA*-infected eyes were highly correlated with each other, whereas all WT-infected and one WT+OxPAPC eye were highly correlated with one another. On the Y-axis, we grouped and color-coded these inflammatory genes based on their function: chemoattractants, complement, innate immune genes, cytokines, kinases, and peptidases. Together, the distinct gene expression clusters and expression patterns demonstrate that mouse ocular inflammatory gene expression was muted if SLP was absent in the infecting strain or if infected eyes were treated with the TLR inhibitor OxPAPC.

### Interfering with TLR2 and TLR4 activation reduced ocular inflammatory responses

We further probed our NanoString data to quantify genes that were altered in our experimental groups. Figure 2A-C and 2G i-ii demonstrate that among the 248 mouse inflammatory genes, 137 genes were significantly upregulated in WT-infected eyes compared to uninfected eyes. However, the number of significantly upregulated genes in WT+OxPAPC eyes and Δ*slpA*-infected eyes was only 44 and 34, respectively (Figure 2G i) compared to that of uninfected eyes. Compared to uninfected eyes, expression of 9 genes, which were common to all non-control groups, was significantly increased (Figure 2G i). Similarly, expression of 91 genes which were common to all non-control groups remained unchanged compared to that of uninfected eyes (Figure 2G ii). We found 66 and 123 genes significantly decreased in WT+OxPAPC and Δ*slpA-*infected eyes, respectively, compared to WT-infected eyes (Figure 2D, E, and G iii). Compared to TLR2 and TLR4 expression in WT-infected eyes, the expression of TLR2 (blue) and TLR4 (red) in WT+OxPAPC and Δ*slpA*-infected eyes was decreased, suggesting that OxPAPC indeed interfered with the innate activation of those two pathways. In Figure 2G iii, the expression of 61 significantly upregulated genes in WT-infected eyes was blunted in WT+OxPAPC and Δ*slpA*-infected eyes. One gene, protein kinase C alpha (PRCKα), was significantly downregulated in WT-infected eyes but upregulated in WT+OxPAPC and Δ*slpA*-infected eyes (Supplementary Table 1). There were no differences in expression of 103 genes in the 4 groups (Figure 2G iv). Compared to uninfected eyes, 55% of total genes were differentially expressed in WT-infected eyes. In contrast, only 18% and 14% of genes were differentially expressed in WT+OxPAPC and Δ*slpA*-infected eyes compared to uninfected eyes, respectively. Compared to WT-infected eyes, expression of 27% and 50% genes were decreased in WT+OxPAPC and Δ*slpA*-infected eyes, respectively (Figure 2F). Taken together, these findings further suggested that inflammation was not as robust when SlpA was absent in *Bacillus* or when TLR2 and TLR4 was inhibited by OxPAPC.

**Figure 2.**
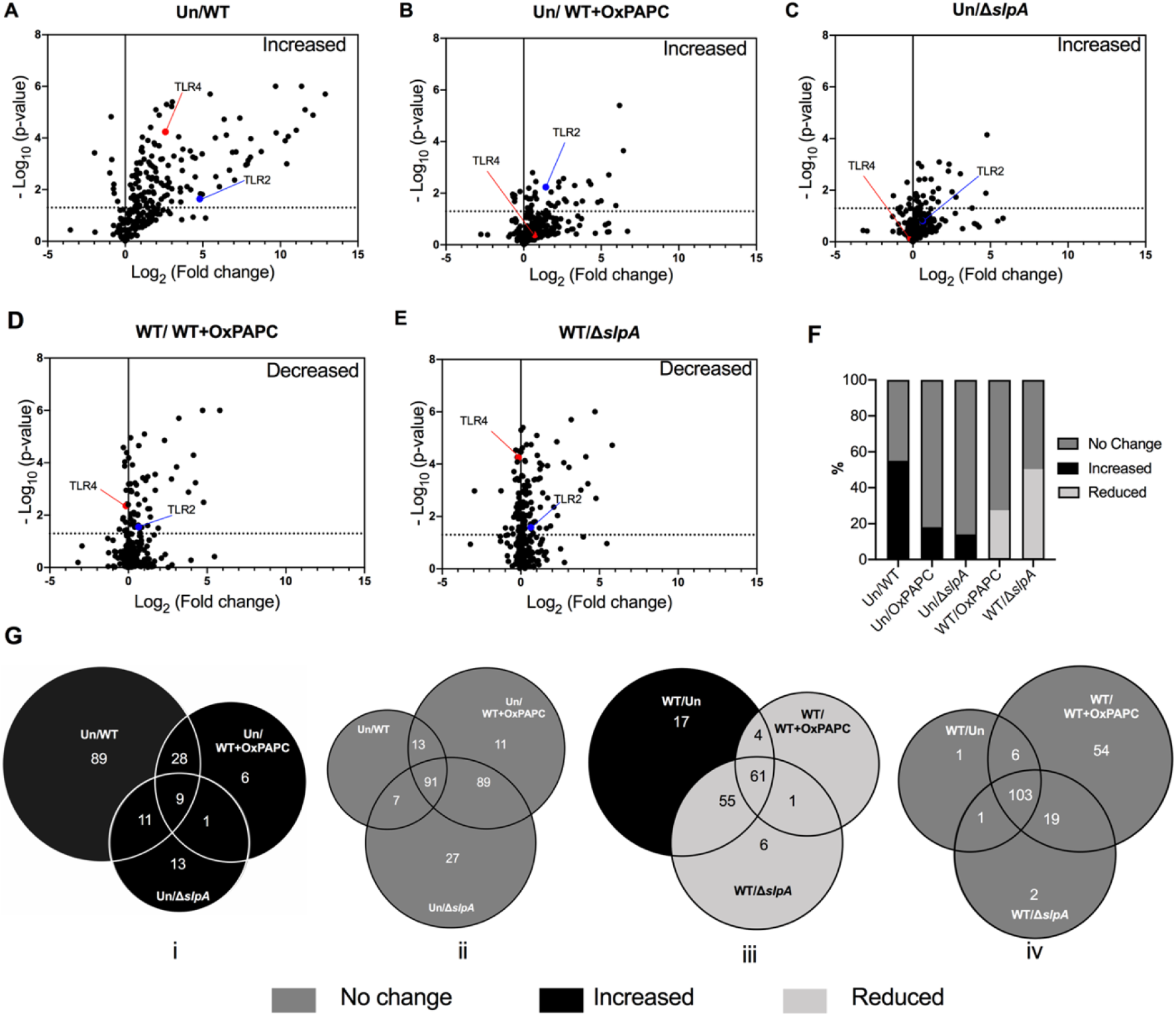
Interfering with TLR2 and TLR4 activation reduced ocular inflammatory responses. Volcano plot analysis of NanoString data derived from WT-infected, WT+OxPAPC, and Δ*slpA*-infected eyes compared with (A-C) uninfected eyes and (D-E) WT-infected eyes. For A-C and D-E the x-axis indicates the log fold change relative to uninfected eyes and WT-infected eyes, respectively. The y-axis shows the negative log10 p-value. Each dot represents an individual gene. TLR2 and TLR4 are indicated by a blue and red dot, respectively. Significance was assessed using multiple t-tests, and p values of < 0.05 (dotted line) were considered to be significant. Any gene above the dotted line was either significantly (A-C) increased or (D-E) decreased. (F) Percent of differentially expressed mouse inflammatory genes in WT+OxPAPC and Δ*slpA*-infected eyes relative to uninfected eyes or WT-infected eyes. (G) Venn diagram (i-iv) showing the number of differentially expressed genes in WT+OxPAPC and Δ*slpA*-infected eyes compared to uninfected or WT-infected eyes. Shading indicates no change in expression (dark gray), increased expression (black), and reduced expression (light gray).

### Interfering with TLR2 and TLR4 activation blunted complement gene expression in the eye

Activation of the complement cascade is one of the first lines of defense in bacterial infection. Studies about the impact of complement factors in the pathogenesis of intraocular diseases is limited. Cobra venom factor-decomplemented Guinea pigs had impaired host defense during intraocular infection with *S. epidermidis* ^52^ and *Pseudomonas aeruginosa* ^53^. Host defenses were restored when complement levels returned to normal in those guinea pigs. In contrast, the absence of complement component C3 did not alter inflammation in a mouse model of experimental *S. aureus* endophthalmitis ^54^. These findings suggest diversity in the contribution of the complement cascade in different endophthalmitis models. Figure 3A depicts the log fold change of complement pathway components in WT-infected, WT+OxPAPC, and Δ*slpA-*infected eyes compared to uninfected eyes. Expression of complement factors b (Cfb), C3, C6, and C7 in WT-infected eyes was 19.6-, 9.9-, 2.9-, and 2.6-fold higher, respectively, compared to uninfected eyes. In contrast, compared to WT-infected eyes, the expression of these genes was decreased in both WT+OxPAPC and Δ*slpA*-infected eyes (Supplementary Table 2). Figure 3B demonstrates the transcript levels of differentially expressed complement factors. We detected increased expression of C3, Cfb, C6, C7, C1q subcomponent-alpha polypeptide (a), C1q subcomponent-beta polypeptide (b), C1r subcomponent A (a), C8 beta polypeptide (b), and C1 s subcomponent (s) transcripts in WT-infected eyes compared to uninfected eyes. Compared to WT-infected eyes, expression of C3, Cfb, and C1ra transcripts were significantly reduced in WT+OxPAPC and Δ*slpA*-infected eyes. Expression of C6, C7, C1qb, and C1s transcripts was also significantly reduced in Δ*slpA*-infected eyes. C8b expression was only reduced in WT+OxPAPC eyes. Expression of hemolytic complement Hc, was increased 1.3-fold in WT-infected eyes compared to uninfected eyes and decreased 1.2-fold and 1.4-fold in WT+OxPAPC and Δ*slpA* infected eyes respectively, compared to WT-infected eyes. Collectively, these findings suggested that *Bacillus* infection-driven complement gene expression in the eye could be prevented by inhibiting TLR2 and TLR4 activation.

**Figure 3.**
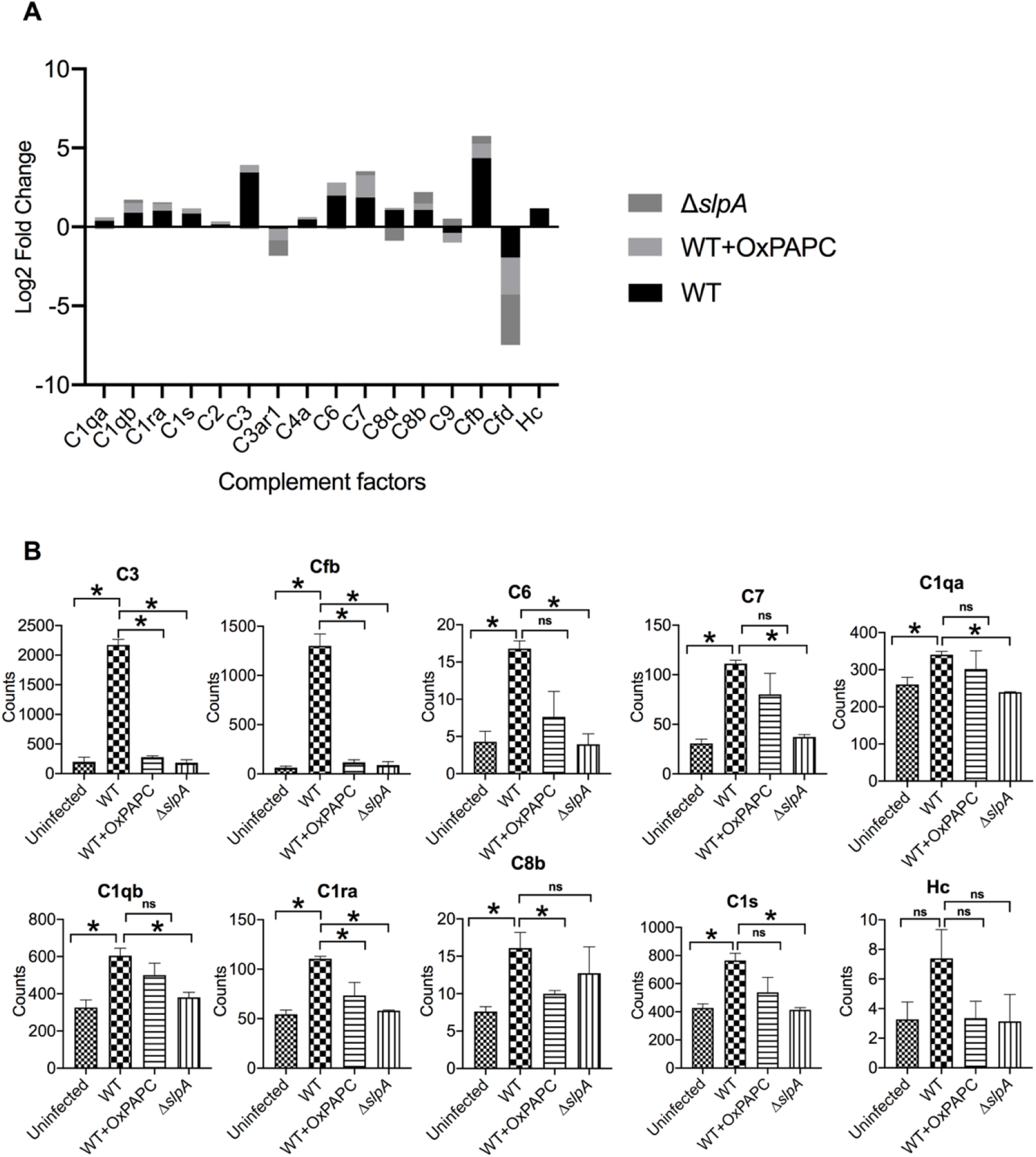
Interfering with TLR2 and TLR4 activation blunted complement gene expression in the eye. Analysis of complement factor expression in WT-infected, WT+OxPAPC, and Δ*slpA*-infected eyes. (A) Fold changes of complement factor expression in WT-infected, WT+OxPAPC, and Δ*slpA-*infected eyes compared to uninfected eyes. (B) Compared to WT-infected eyes, decreased expression of C3, Cfb, C6, C7, C1qa, C1qb, C1ra, C8b, and C1s transcripts were observed in WT+OxPAPC or Δ*slpA*-infected eyes. Values represent the mean±SEM of complement counts at 10 hours postinfection for three different mouse eyes and p-value of < 0.05 was considered significant.

### Interfering with TLR2 and TLR4 activation impacted innate immune gene expression in the eye

Intraocular innate receptors and their adaptors contribute to the pathogenesis of *Bacillus* endophthalmitis and other vision-threatening infections. During *Bacillus* endophthalmitis, severe intraocular inflammation is triggered by TLR2 and TLR4 via MyD88 and TRIF ^36,37,50^. Here, we focused on 41 genes involved in innate immune pathways, which include receptors, adaptors, signaling molecules, and hormones. (Supplementary Table 3). Among the 24 differentially expressed innate immune pathway genes, 11 were highly upregulated in WT-infected eyes compared to uninfected eyes, and were reduced in WT+OxPAPC and Δ*slpA*-infected eyes compared to WT-infected eyes (Figure 4A). Figure 4B depicts a heatmap of the expression of TLRs and other innate receptors. As expected, we observed increased expression of TLR2 and TLR4 transcripts in WT-infected eyes. In addition, we observed elevated expression of TLR6, TLR8, nucleotide-binding oligomerization domain-containing protein (NOD) 2, and LRR- and pyrin domain-containing protein (Nlrp3) in WT-infected eyes. Expression of these innate genes was significantly decreased in WT+OxPAPC or Δ*slpA*-infected eyes compared to WT-infected eyes. The expression of other innate immune genes, such as MyD88, NF-κB, v-rel reticuloendotheliosis viral oncogene homolog (Rel) A, and v-rel oncogene related (Rel) B, prostaglandin-endoperoxide synthase (Ptsg) 2, signal transducer and activator of transcription (STAT) 2 and 3, and heat shock protein (Hspb) 1 was significantly increased in WT-infected eyes compared to uninfected eyes (Figure 4C). Expression of these genes was decreased in WT+OxPAPC and Δ*slpA*-infected eyes compared to WT-infected eyes. Among the 41 innate immune genes we analyzed, expression of Ptgs2, Nlrp3, TLR2, TLR6, and NOD2 were 218-, 140-, 26.8-, 19.8-, and 17.5-fold greater than in uninfected eyes and were the top five upregulated innate immune genes in WT-infected eyes. We observed a negative fold change of these genes in WT+OxPAPC and Δ*slpA*-infected eyes compared to WT-infected eyes (Supplementary Table 3). Here, we identified several innate immune genes whose expression was not changed at 4 hours postinfection in *Bacillus* endophthalmitis. Furthermore, we observed dampened expression of those innate immune genes when TLR2 and TLR4 activation did not occur. These results suggested that the innate immune gene expression in *Bacillus* endophthalmitis occurring during robust inflammation could be prevented by inhibiting TLR2 and TLR4 activation.

**Figure 4.**
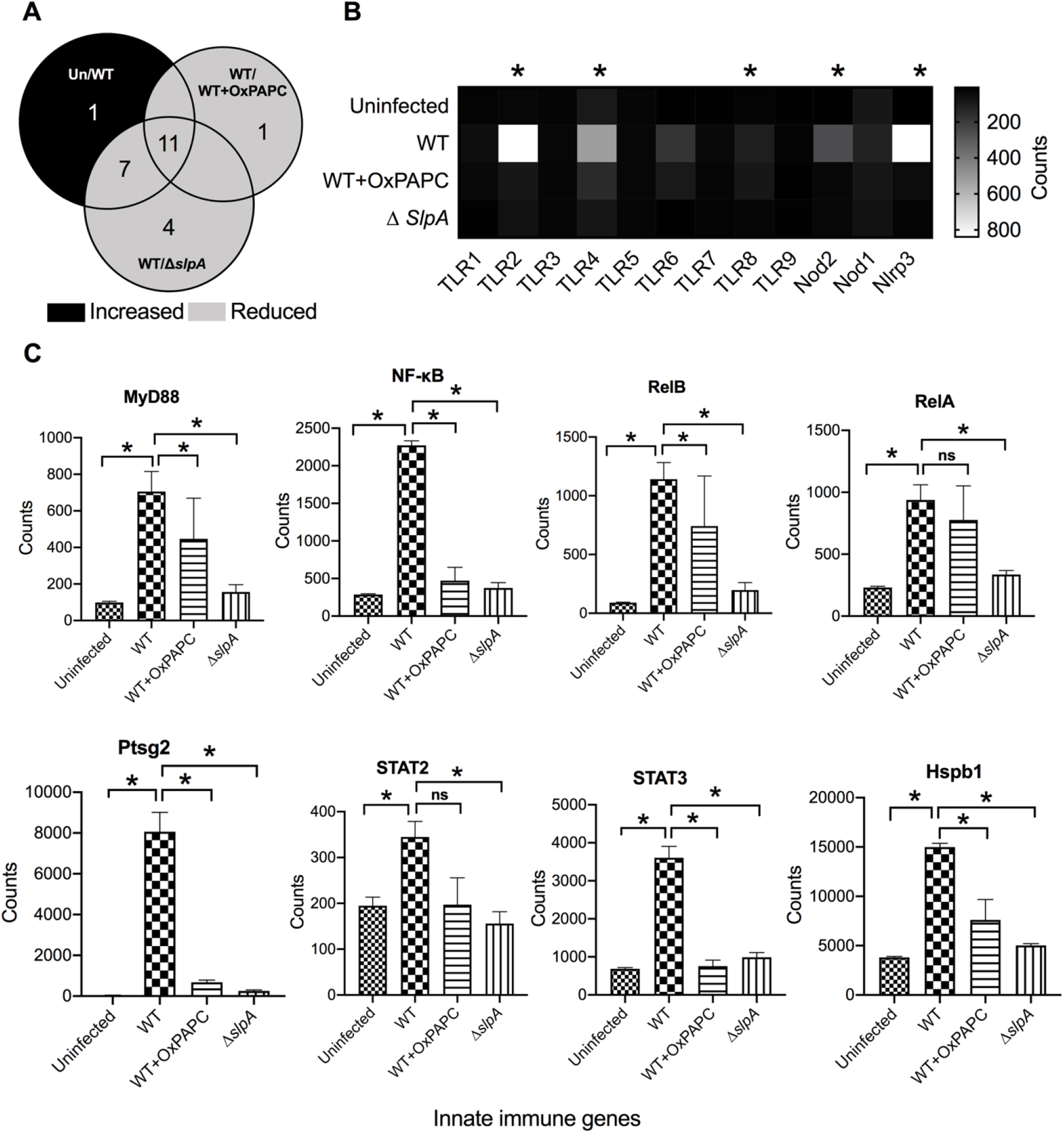
Interfering with TLR2 and TLR4 activation impacted innate immune gene expression in the eye. Expression of innate immune genes in mouse eyes. (A) Venn diagram showing the number of innate immune genes differentially upregulated between the groups. (B) Heatmap showing the expression of extracellular and intracellular innate receptors. TLR2, TLR4, TLR6, TLR8, intracellular receptor Nod2, and Nlrp3 were significantly upregulated in WT untreated eyes compared to uninfected eyes. (C) Innate immune genes such as MyD88, NF-κB, RelA, RelB, Ptsg2, Hspb1, STAT2, STAT3, and Hspb1 were significantly upregulated in WT-infected eyes compared to uninfected eyes. Values represent the mean±SEM of transcript counts at 10 hours postinfection for 3 different mouse eyes and *P< 0.05 was considered significant.

### Interfering with TLR2 and TLR4 activation reduced cytokine gene expression in the eye

During endophthalmitis, cytokines and chemokines drive ocular inflammation. In TNFα^−/−^ and CXCL1^−/−^ mice, neutrophil recruitment was suppressed. However, this resulted in an increase in bacterial load and disease severity in TNFα^−/−^ mice^44^, but no change in bacterial load and a reduction in disease severity in CXCL1^−/−^ mice^45^. These contrasting outcomes suggest a potential role of additional inflammatory cytokines during infection. Here, we focused on the expression of 51 mouse inflammatory cytokines during *Bacillus* endophthalmitis. Figure 5A depicts the log-fold changes of these genes in our experimental groups. Eighteen inflammatory cytokine genes were significantly increased in WT-infected eyes compared to uninfected eyes. These genes were decreased in WT+OxPAPC or Δ*slpA*-infected eyes compared to WT-infected eyes (Figure 5B, Supplementary Table 4). Figure 5C depicts the expression of IL-6, TNFα, and IL-1β and several other pro-inflammatory cytokines in our infection and treatment groups. In addition to IL-6, TNFα, and IL-1β, the expression of colony-stimulating factor (CSF)3, CSF2, IL-1α, IL-23α, transforming growth factor (TGF)β1, and IL-12β were significantly increased WT-infected eyes compared to uninfected eyes. Expression of CSF3, IL-6, IL-1β, CSF2, IL-1α, TNFα, and IL-23α was significantly decreased in both WT+OxPAPC and Δ*slpA*-infected eyes compared to WT-infected eyes. Expression of TGFβ1 and IL-12β was reduced only in Δ*slpA*-infected eyes compared to WT-infected eyes.

**Figure 5.**
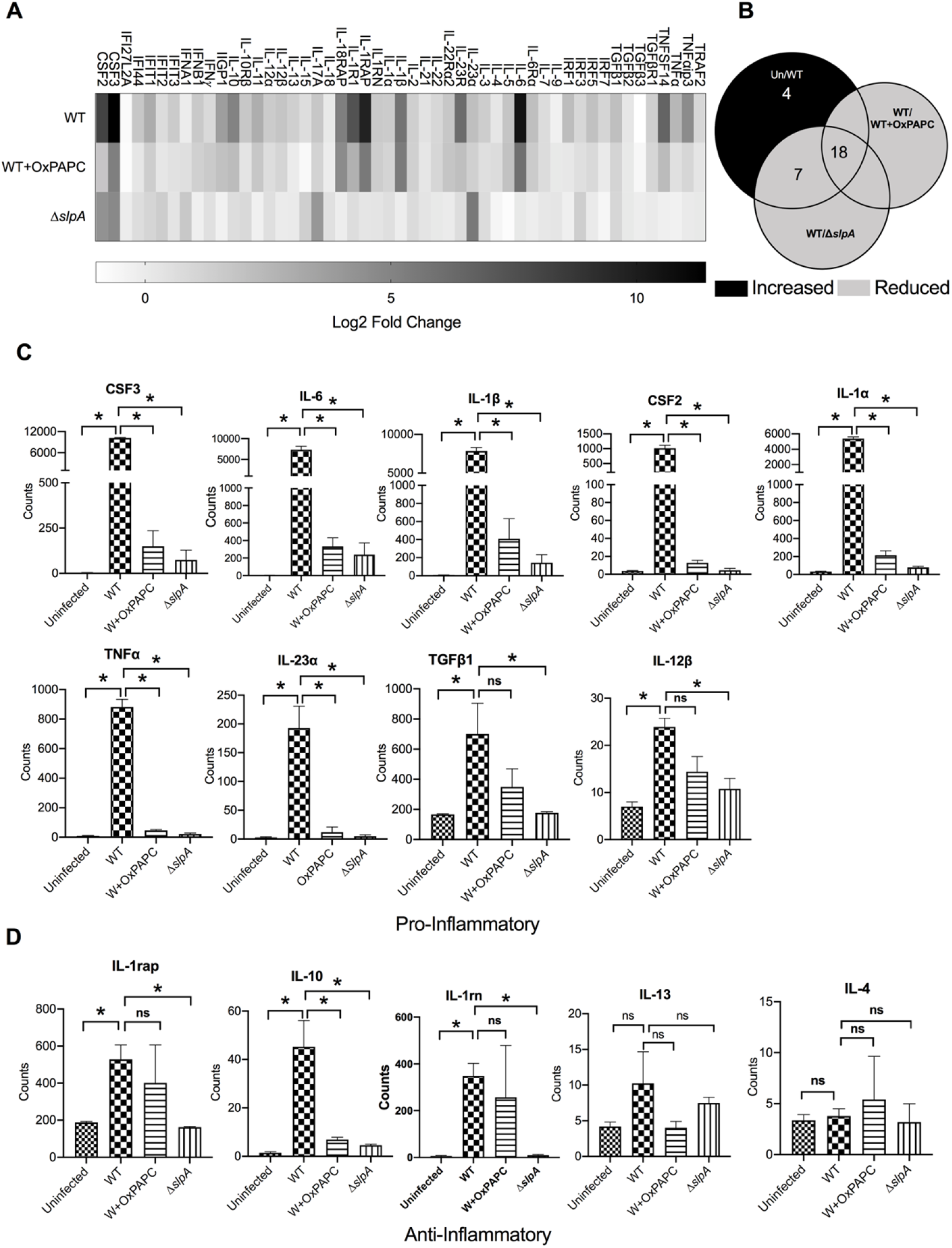
Interfering with TLR2 and TLR4 activation reduced cytokine gene expression in the eye. Expression of inflammatory cytokines from WT-infected, WT+OxPAPC, Δ*slpA-*infected eyes were analyzed. (A) Log fold change of 51 mouse inflammatory cytokines from WT-infected, WT+OxPAPC, and Δ*slpA-*infected eyes, compared to uninfected eyes. The darkest shading corresponds to the greatest fold changes.(B) Venn diagram showing the number of cytokine genes differentially expressed between the groups. (C) Elevated expression of pro-inflammatory cytokines CSF3, IL-6, IL-1β, CSF2, IL-1α, TNFα, IL-23α, TGFβ1, and IL-12β in WT-infected eyes compared to uninfected eyes. Compared to WT-infected eyes, expression of these pro-inflammatory cytokines was significantly reduced in WT+OxPAPC or Δ*slpA*-infected eyes. (D) Compared to uninfected eyes, expression of anti-inflammatory cytokines IL-1rap, IL-10, and IL-1rn was elevated in WT-infected eyes, but was reduced significantly in WT+OxPAPC or Δ*slpA*-infected eyes. Values represent the mean±SEM of transcript counts at 10 hours post infection (N=3). *P< 0.05 was considered significant.

Pro-inflammatory cytokines act in concert with specific cytokine inhibitors to regulate the human immune response. The expression of anti-inflammatory cytokines IL-1 receptor accessory protein (rap), IL-10, and IL-1 receptor antagonist (rn) was significantly increased in WT-infected eyes compared to uninfected eyes, but significantly reduced in Δ*slpA*-infected eyes compared to WT-infected eyes. The expression of IL-10 was reduced only in WT+OxPAPC eyes compared to WT-infected eyes (Figure 5D). The expression of anti-inflammatory cytokines IL-13 and IL-4 were not changed in all groups. Relative to uninfected eyes, expression of CSF3, IL-6, IL-1β, CSF2, IL-1α, and TNFα was 50-fold or higher (Supplementary Table 4). These findings not only identified several inflammatory cytokines which were not expressed at 4 hours postinfection in *Bacillus* endophthalmitis, but also identified blunted expression of others when TLR2 and TLR4 activation was blocked. Collectively, these findings suggested that the innate inhibition could be a viable strategy to avert inflammatory cytokine production during *Bacillus* endophthalmitis.

### Interfering with TLR2 and TLR4 activation reduced chemoattractant gene expression in the eye

A hallmark of *Bacillus* endophthalmitis is robust influx of inflammatory cells into the posterior chamber of the eye. Chemokines act as chemoattractants to guide the migration of these cells. Figure 6A depicts the log fold changes of 27 inflammatory chemokines and receptors that belong to CXC and CC groups. Fifteen chemokines were significantly increased in WT-infected eyes compared to uninfected eyes. Compared to WT-infected eyes, expression of these 15 chemokines was significantly reduced in WT+OxPAPC or Δ*slpA*-infected eyes (Figure 6B, Supplementary Table 5). Figure 6C depicts the expression of chemoattractants in our infection and treatment groups. Expression of CXCL1, CXCL2, CXCL10, CCL2, CCL3, CXCL3, CCL20, and CCL4 was elevated in WT-infected eyes compared to uninfected eyes. Expression of these genes was reduced 10-fold or greater in WT+OxPAPC or Δ*slpA*-infected eyes compared to WT-infected eyes. Figure 6D depicts analysis of the expression of chemokine receptors. Expression of chemokine (C-C motif) receptor (CCR)1, chemokine (C-X-C motif) receptor (CXCR) 2, CXCR4, and CCR7 was greater in WT-infected eyes compared to uninfected eyes, but was significantly reduced in WT+OxPAPC or Δ*slpA*-infected eyes compared to WT-infected eyes. CCR7 expression was significantly increased in WT-infected eyes compared to uninfected eyes, and significantly reduced in Δ*slpA*-infected eyes compared to WT-infected eyes. There was no change in CCR7 expression between WT-infected and WT+OxPAPC eyes. Expression of CXCL2, CXCL1, CXCL3, CCL20, CCL4, CCL3, and CCL2 was 500-fold or higher relative to that of uninfected eyes (Supplementary Table 5). In addition to confirming our retina-specific transcriptome data at 4 hours postinfection with *Bacillus* ^50^, these findings identified the expression of additional chemoattractants in the whole eye during the later stages of infection. Altogether, these results demonstrated that the chemokine response is blunted when SlpA is absent in *Bacillus* and when TLR2 and TLR4 are inhibited by OxPAPC. Collectively, these results suggested the inhibition of TLR2 and TLR4 pathways might be a viable strategy to block the chemoattractant synthesis during *Bacillus* endophthalmitis, arresting the influx of inflammatory cells into the eye.

**Figure 6.**
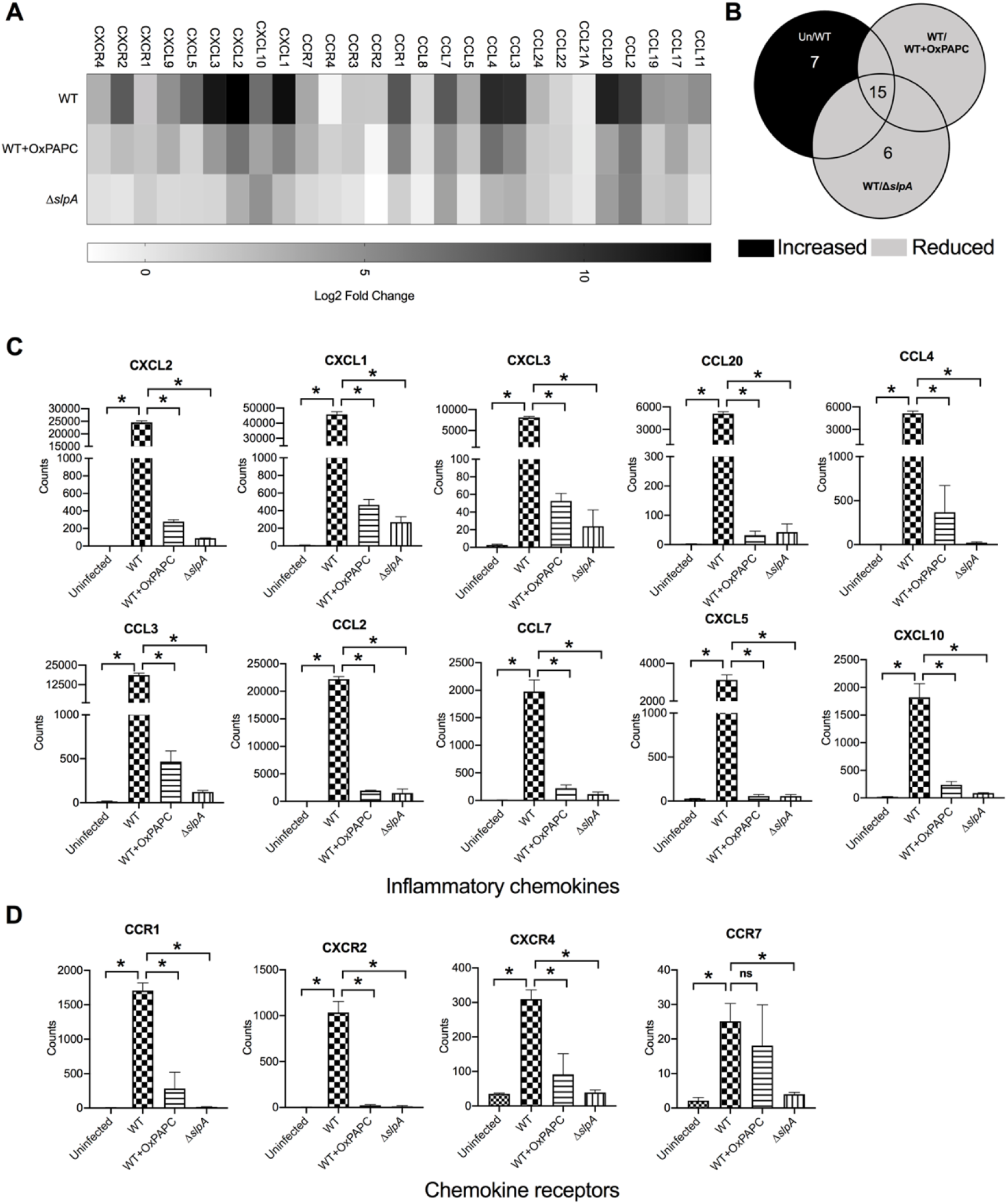
Interfering with TLR2 and TLR4 activation reduced chemoattractant gene expression in the eye. NanoString analysis of CXC and CC chemokine groups analyzed in WT-infected, WT+OxPAPC, and Δ*slpA*-infected mouse eyes. (A) Log fold changes of inflammatory chemokines from WT-infected, WT+OxPAPC, and Δ*slpA-*infected eyes, compared to uninfected eyes. The darker shading corresponds to the greatest log-fold changes.(B) Venn diagram showing the number of chemoattractants differentially expressed between the groups. (C) At 10 hours postinfection, expression of CXCL2, CXCL1, CXCL3, CCL20, CCL4, CCL3, CCL2, CCL7, CXCL10, and CXCL5 were upregulated in WT-infected, and significantly reduced in WT+OxPAPC and Δ*slpA*-infected eyes. (D) Expression of chemokine receptor CCR1, CXCR2, CXCR4, and CCR7 was also significantly upregulated in WT-infected eyes and reduced in WT+OxPAPC and Δ*slpA*-infected eyes.Values represent the mean±SEM of transcript counts at 10 hours postinfection for three individual mouse eyes. *P< 0.05 was considered significant.

## DISCUSSION

The inflammatory response is generally defined as a protective reaction to stimulation of the host defense system to invading pathogens or endogenous signals. When this protective response goes unchecked, dysregulated inflammatory responses occur and can worsen the disease severity. Cells involved in immune responses produce plethora of mediators including cytokines, chemokines, antibodies, and complement to aid in the war against invading pathogens. Although inflammatory responses protect tissue against these insults, the inflammation itself could cause damage. Immune privileged tissues resist immunogenic inflammation through multiple mechanisms. A well-controlled system to regulate the inflammatory response in the eye is necessary ^35^.

We reported that *Bacillus* reach their maximum growth faster than most other Gram-positive endophthalmitis pathogen in the eye ^9^*. Bacillus* endophthalmitis is well recognized for creating an aggressive acute inflammatory response that results in retinal damage with rapid loss of vision within 12-24 hours despite antibiotic and anti-inflammatory treatment. Avirulent *S. epidermidis* is generally cleared by an active but not overly robust inflammatory response. Current therapies for this severe blinding infection include intravitreal antibiotics that can sterilize the eye if administered at an early phase of infection ^1–8^. The use of corticosteroids for anti-inflammatory therapy in endophthalmitis is common, but their efficacy in this disease is controversial ^55–61^. Therefore, identifying critical elements of inflammatory pathways that can be targeted to inhibit the intraocular inflammation are viable prospects for future therapeutics in *Bacillus* endophthalmitis.

One of the earliest steps of any inflammatory response is the recognition of the invading organisms by the host innate defense system. The retina is comprised of different types of cells necessary for the visual cycle, maintaining retinal integrity, and homeostasis. These cells also act as a defensive barrier to protect against invading pathogens by expressing innate receptors. We reported that during infection, *Bacillus* migrates in close proximity to the inner limiting membrane (ILM) of the retina ^11^, facilitating potential interactions between the retina and invading microbes. As such, the outer most layer of the bacterium may be the first to interact with innate receptors on cells lining the ILM.

S-layer protein (SLP), the outer most layer of the *Bacillus* envelope, serves as a protective 444 barrier for the pathogen. Besides maintaining cell wall integrity, SLP provides additional benefits for the microbes to survive in diverse host environments, but can also activate the immune system ^26,62,63^. We recently reported that the absence of *Bacillus* SLP reduced disease severity and damage to the eye in murine experimental endophthalmitis ^33^. Activation of innate immunity and its effects on host tissue play a crucial role in the severity and poor visual outcomes of *Bacillus* endophthalmitis. In a follow-up study, we reported *Bacillus* SLP as an activator of the TLR2 and TLR4 pathways ^34^. In a recent transcriptome study of retinas of eyes infected with *Bacillus*, we reported elevated expression of 72 genes at 4 hours postinfection and decreased expression of these 72 genes in the absence of TLR4 ^50^. The reduced inflammation and improved clinical outcomes in the absence of TLR2 ^36^ or TLR4 ^37^ or following TLR2 and TLR4 inhibition ^34^ suggests innate immune pathway inhibition as a prospective option for the anti-inflammatory arm of treatment of *Bacillus* intraocular infection.

The nature of ocular tissue damage arising from an excessive inflammatory response in bacterial endophthalmitis is not completely understood. Based on our recent studies demonstrating positive clinical outcomes when TLR2 and TLR4 pathways are blocked, we further probed this treatment strategy by using NanoString to compare gene expression in whole eyes in our experimental groups. NanoString is better for targeted transcriptomics since there is no need for preparing gene libraries, enzymes, and processing as in other next generation sequencing (NGS) techniques ^64^. Unlike other alternatives, NanoString does not require polymerase activity, and therefore the chance of introducing bias is lower ^65–67^. NanoString can identify RNAs in a heterogeneous sample, even one that contains cells from different species. In this study, we focused on 248 mouse genes potentially associated with the inflammatory response in experimental murine *Bacillus* endophthalmitis. Distinct clusters of infection groups in our principal component analysis suggested a similar pattern of gene expression within each group. Except for one eye in the WT+OxPAPC group, we observed specific clusters in our groups. One explanation for this outlier could be a microinjection error during intraocular OxPAPC delivery. Overall, however, the clustering of uninfected, Δ*slpA*-infected, and WT+OxPAPC eyes suggested transcriptional similarities among these groups.

Most of the genes analyzed were differentially expressed in WT-infected eyes compared to uninfected eyes. Expression of most of these genes were reduced in WT+OxPAPC, and Δ*slpA*-infected eyes compared to WT-infected eyes. Expression of 61 genes which were significantly increased in WT-infected eyes compared to uninfected eyes were decreased in WT+OxPAPC or Δ*slpA*-infected eyes compared to WT-infected eyes (Supplementary Table 1). Only one gene, protein kinase C alpha (PRKCα), was decreased in WT-infected eyes compared to uninfected eyes but significantly increased in Δ*slpA*-infected and WT+OxPAPC eyes. PRKCα is an essential modulator of rod photoceptor and bipolar cell function. PRKCα and few other isoforms of PRKC are reported to be present in ganglion, amacrine, and RPE cells. PRKCα has a role in a variety of cellular functions, including proliferation, apoptosis, differentiation, motility, and inflammation ^68–71^. It has been reported that TLR1/2-driven activation of MAPK, NF-kB, and AP-1, as well as secretion of TNF-α, IL-6, and IL-10 by dendritic cells requires the activation of PRKCα ^70^. Overexpression of PRKCα resulted in loss of tight junctions, which reduced the transepithelial permeability in an epithelial cell line LLC-PK1 ^72^. Nitric oxide is an effector molecule of innate immunity and inflammatory modulator in different inflammatory conditions, and PRKCα has been linked to the regulation of nitic oxide production in vascular smooth muscle, murine macrophages, and murine microglia ^73^. Activation of protein kinase C (PKC) induced the production of various inflammatory cytokines from human bronchial epithelia cells, suggesting that PKC activation might be crucial for the initial defense mechanism in the airway against pathogens ^74^. The absence of PRKC isoform δ in mice resulted in reduced inflammatory mediator production and neutrophil infiltration in an asbestos-associated disease model ^75^. We observed decreased expression of PRKCα in WT-infected eyes compared to uninfected eyes, which suggested a potential anti-inflammatory property of PRKCα. However, this was not reflected in the pathogenesis *in vivo*, since we found increased PMN influx in WT-infected eyes. PKC is expressed differentially in various mammalian tissues, and is highly enriched in brain and lymphoid organs ^76^. Together, this suggests that the role of PRKC in inflammation may be tissue and organ-specific.

Improving treatment options is a main focus of our research in endophthalmitis. This is especially critical for *Bacillus* endophthalmitis in which the majority of infected eyes lose significant vision. The contribution of the host immune response in mediating bystander damage and visual function loss is still not well understood. Complement is an integral component of the innate host defense against infection and a prime mediator of tissue inflammation. We recently reported that pentraxin 3 (PTX3), which activates the classical complement pathway via C1q to facilitate pathogen recognition and clearance, was expressed as early as 4 hours in retinas of mouse eyes infected with *Bacillus* ^50^. In this study, TLR2 and TLR4 interference resulted in decreased expression of PTX3 and other complement pathway-related genes. In endophthalmitis caused by *S. epidermidis* and *Pseudomonas aeruginosa*, complement exhibited a beneficial effect in ocular defense ^52,53^. However, for *S. aureus* endophthalmitis, complement was not protective ^54^. Since the interplay between complement and bacteria is relevant in inflammation, studying the role of complement pathway-associated genes in the pathogenesis of a rapidly blinding infection like *Bacillus* endophthalmitis may be valuable.

Innate recognition activation of associated pathways are complex events requiring coordinate regulation of multiple cellular signaling components. We previously reported the role of TLR2 and TLR4 in the pathogenesis of *Bacillus* endophthalmitis ^36,37^. TLR2 is also important for inflammatory responses in *S. aureus* endophthalmitis ^77^. TLR6 forms a heterodimer with TLR2 and recognizes diacylated lipopeptides such as lipoteichoic acid on the cell wall of Gram-positive bacteria. Synergistic interactions of TLR2/6 and TLR9 provide resistance against lung infection ^78,79^. TLR8, which is an innate endosomal receptor, usually senses viral infection ^80^. However, TLR8 may also detect RNA from *S. aureus*, which occurs when the bacterium is internalized and degraded inside a phagocytic cells. Activation of TLR2 could serve as an inhibitor of TLR8 activation inside the cell ^81^. This counter-inhibition might serve as a safety mechanism to avoid exaggerated immune activation. Since TLR2 and TLR8 sense different PAMPs, this cross-regulation represents a fine-tuning to the excessive immune response towards different classes of pathogens. Neither TLR6 nor TLR8 expression was detected at 4 hours postinfection in retinas of *B. cereus*-infected mice^50^. TLR6 and TLR8 were expressed, but not to a great degree at 10 hours in *B. cereus*-infected mouse eyes in this study, so this expression may emanate from tissues or cells other than in the retina. Inflammasomes are critical as they protect the tissue against pathogens and maintain homeostasis with commensal microbes ^82,83^. While we reported that the absence of Nlrp3 in Nlrp3^−/−^ global knockout mice did not affect the clinical outcomes in *Bacillus* endophthalmitis at 8 hours postinfection ^61^, a recent report suggested that non-hemolytic enterotoxin (NHE) from *B. cereus* activated the NLRP3 inflammasome and pyroptosis in primary BMDMs, and that NHE and hemolysin BL functioned synergistically to induce inflammation in mice ^84^. Here, we observed significant increased expression of TLR2, TLR4, and Nlrp3 in WT-infected eyes compared to uninfected eyes. Expression of these innate receptors were reduced after TLR2 and TLR4 inhibition or infection with the Δ*slpA* mutant. These results suggested that interference of innate pathways affected the expression of other innate receptors in the eye, perhaps further limiting inflammation.

An absence of MyD88 or TRIF in mice resulted in improved clinical outcomes of *Bacillus* endophthalmitis ^38^. The importance of MyD88 has also been reported for *S. aureus* endophthalmitis ^85^. Lipid mediators derived from arachidonic acid control cell proliferation, apoptosis, metabolism, and migration ^86^. Prostaglandin-endoperoxide synthase (Ptgs) 2 converts arachidonic acid to prostaglandin endoperoxide H2 (PGE2), which has an immunomodulatory effect and serves to prevent the activation of neutrophils ^87^. Ptgs2 was upregulated in a TLR4-dependent manner in the retinas of mice infected with *Bacillus* ^50^. Activation of TLRs and their adaptors transfer the activation signal via a series of factors that ultimately activate inflammatory transcription factor NF-κB in the nucleus. We recently reported that *Bacillus* SLP activated NF-κB in human retinal Muller cells ^33^. Here, we observed increased expression of MyD88, Ptgs2, RelA, RelB, STAT2, STAT3, and Hspb1 in WT-infected eyes compared to that of uninfected eyes. We also observed blunted expression of these innate immune genes when TLR2 and TLR4 pathways were interfered with. Unfortunately, TRIF expression was not included in the NanoString panel, so this was not tested. Phosphorylation of STAT3 and Hspb1 influences NF-kB activation and promotes the inflammatory response. Retinal transcriptome analysis of *S. aureus* endophthalmitis showed elevated expression of STAT3 in a TLR2-dependent manner ^88^. SOCS3, which is a negative regulator of cytokine signaling, inhibits STAT3 phosphorylation and prevents typically excessive activation of proinflammatory genes ^89^. The enhanced expression of STAT3 in our study may help to explain why increased TLR4-dependent expression of SOCS3 failed to negatively regulate inflammatory gene expression.

The *Bacillus* cell wall is highly inflammogenic. *Bacillus* SLP triggers the expression of inflammatory mediators CXCL1, TNFα, CCL2, and IL-6 in human retinal Muller cells ^33^. Inflammatory cytokines, which are polypeptides, can act as intercellular messengers and mediate the process of inflammation and repair. In the eye, in response to an infection or injury, inflammatory cytokines can be produced by corneal epithelium, RPE, microglia, macrophages, endothelial cells, and other immune cells ^90^. Here, we analyzed the expression of 51 inflammatory cytokines and found 50-fold or greater expression of IL-1α, IL-1β, IL-6, CSF2, CSF3, and TNFα in WT-infected eyes (Supplementary Table 4). CSF is a glycoprotein that stimulates the production of neutrophil granulocytes and their release into the blood and the survival, proliferation, differentiation, and function of neutrophil precursors and mature neutrophils ^91,92^. Quite a bit is known about CSF’s role in treating neutropenia and hematopoietic mobilization, but information about its role in ocular infection is limited. Neutrophils enter the eye as early as 4 hours postinfection and are the primary immune cells that migrate into the vitreous after infection ^43^. Because of its role in neutrophil migration, the increased expression of CSF in the eye may contribute to the rapid migration of neutrophils into the eye in experimental *Bacillus* endophthalmitis. Because we did not observe an increase in the expression of CSF at 4 hours postinfection in retinas of infected mouse eyes ^50^, the source of CSF may not be the retinal cells at this early stage of infection.

IL-6 is both a pro-inflammatory and anti-inflammatory cytokine, depending upon the environment ^93^. IL-6 expression increases during the course of bacterial endophthalmitis ^43,77^ in a TLR2- and TLR4-dependent manner ^36,37^. Expression of IL-6 and IL-1β was also increased with time in retinas of *S. aureus*-infected mice; however, compared to wild type, expression of these mediators was reduced when TLR2 was absent ^88^. Surprisingly, the absence of IL-6 in IL-6^−/−^ mice did not affect the overall outcome of *Bacillus* endophthalmitis ^45^. Here, interference with TLR 2 and TLR4 also resulted in reduced expression of the anti-inflammatory cytokine IL-10 at 10 hours postinfection. Intravenous administration of IL-10, a potent inhibitor of cytokine production, reduced neutrophil chemotaxis in LPS-induced uveitis ^94,95^. Our results suggest that the expression of anti-inflammatory cytokines may not affect the clinical outcome in our model because the expression of anti-inflammatory cytokines in WT-infected eyes did not affect the expression of IL-6, TNF-α, or IL-1β.

Chemokines are inflammatory mediators that serve as chemoattractants and are divided into two significant subfamilies (CXC and CC chemokines) based on the spacing of the first pair of N-terminal cysteine residues ^96–98^. The CXC chemokines usually recruit neutrophils, whereas CC chemokines tend to attract monocytes and involve in the pathogenesis of immune mediated inflammation ^96–98^. Activation of TLR2 on retinal microglia ^99^, and Muller glia ^100,101^ by *S. aureus* resulted in the synthesis of CXC chemoattractants. Among the 248 mouse inflammatory genes analyzed, CXCL2, CXCL1, and CXCL3 were the most highly expressed chemokines in WT-infected eyes (Supplementary Table 5). CXCL2, CXCL1, and CXCL3 are highly homologous and are powerful neutrophil chemoattractants ^96–98^. We also observed increased expression of the CC chemokines CCL2/MCP-1, CCL3/macrophage inflammatory protein (MIP)-1α, CCL4/MIP-1β, CCL7/monocyte-chemotactic protein 3 (MCP3), and CCL20/Macrophage Inflammatory Protein-3 (MIP3A) in WT-infected eyes. CCL2 does not function as a chemoattractant for neutrophils, but does recruit basophils, monocytes, memory T-cells, and dendritic cells to the sites of inflammation induced by either tissue injury or microbial infection ^102^. CCL2 expression in Muller cells regulate the infiltration of monocytes and microglia following retinal injury ^102^. CCL2 has been associated with acute inflammatory conditions ^98^, and recruits and activates neutrophils by binding to CCR1, CCR4, and CCR5 ^102,103^. CCL20 strongly attracts lymphocytes and weakly attracts neutrophils ^104^. Expression of CCL2 and CCL20 can be induced by microbial factors such as lipopolysaccharides and inflammatory cytokines such as IL-6 and TNFα ^104–108^. We reported TLR4-dependent expression of CCL2 and CCL3 in the retinas of *Bacillus*-infected mouse eyes at 4 hours postinfection ^50^. Here, we demonstrated blunted expression of CCL2, CCL3, CCL7, CCL4, and CCL20 when TLR2 and TLR4 activation was interfered with. CXCR2, CXCR4, CCR1 are the major CXC and CC chemokine receptors that bind and respond to chemokines to mobilize immune cells. CXCR1 and CXCR2 interact with CXCL1-8 and are expressed on the neutrophil surface ^103^. Therefore, the increased expression of CXC and CC receptors and chemoattractants in WT-infected eyes may be related to the presence of neutrophils in the intraocular environment at this stage of infection. We reported that the absence of CXCL1 in CXCL1^−/−^ mice or intraocular injection of anti-CXCL1 improved the clinical outcome of *Bacillus* endophthalmitis, resulting in minimal inflammation and retained retinal function ^45^. The absence of TLR2 resulted in reduced expression of CXCL2 in mouse retinas during experimental *S. aureus* endophthalmitis ^88^. Although CCL2 is not known to recruit neutrophils, treatment with anti-CCL2 or anti-CCL3 significantly reduced neutrophil infiltration into the cornea, resulting in decreased corneal damage in a mouse model of *Pseudomonas aeruginosa* keratitis ^109^. The expression of more than half of the chemokines that we analyzed was blunted when TLR2 and TLR4 activation was interfered with (Supplementary Table 5). Together, these findings suggested a therapeutic potential for targeting CC and CXC chemokines in controlling ocular inflammation during *Bacillus* endophthalmitis, and possibly other forms of endophthalmitis.

During ocular infection with an avirulent organism, the innate immune response is usually sufficient to clear the infection. However, ocular infections caused by a more virulent pathogen such as *Bacillus* are not easily cleared, and the robust innate response induced by *Bacillus* cell wall components can lead to significant host-mediated damage. Therapeutics designed to counteract the activities of individual bacterial products might not prevent the damage caused by numerous other toxic microbial factors, nor reduce inflammation-induced injury. The fact that the absence of *Bacillus* S-layer had such a profound effect on intraocular inflammation and infection highlights S-layer as an important virulence factor in this disease. The ability of *Bacillus* S-layer to activate both TLR2 and TLR4 may partially explain why *Bacillus* endophthalmitis results in such an explosive inflammatory response.

Innate inhibition by interfering the TLR pathways has been promising in several inflammation-associated diseases such as Gram-negative bacterial sepsis, acute lung inflammation injury, atherosclerosis, acute and chronic inflammatory pain ^47,49,110–115^. The negative regulation of inflammation by oxidized phospholipid has been effective for the treatment of these diseases. Oxidized phospholipids have been shown to inhibit LPS-induced pyroptosis, IL-1β release, and septic shock, providing a basis for therapeutic innate immune interference in Gram-negative bacterial sepsis ^47^. Oxidized phospholipids also markedly attenuated LPS-induced tissue inflammation, barrier disruption, and cytokine production in rats ^111^. Since the inflammatory response triggered by the innate immune response to *Bacillus* endophthalmitis may cause severe damage to ocular tissues, targeting these pathways could be a viable anti-inflammatory approach which also preserves retinal function. We identified 25 inflammatory genes (Table 1) which were upregulated 50-fold or higher after infection with WT *Bacillus*, and demonstrated reduced expression of these 25 inflammatory genes after OxPAPC-mediated TLR2 and TLR4 inhibition or infection with a SLP-deficient *Bacillus*. The expression of 12 chemoattractants among these 25 genes was likely the driver of excessive neutrophil infiltration during *Bacillus* infection. Together, these results suggest that *Bacillus* SLP potentially triggered the expression of these inflammatory genes, which could be prevented by inhibiting the activation of TLR2 and TLR4 pathways. Although we demonstrated improved clinical outcomes in *Bacillus* endophthalmitis when infected eyes were treated with OxPAPC, we did not test co-administration with antibiotics. The administration of anti-inflammatory drugs alone is not an acceptable standard of care for endophthalmitis.

**Table 1.**
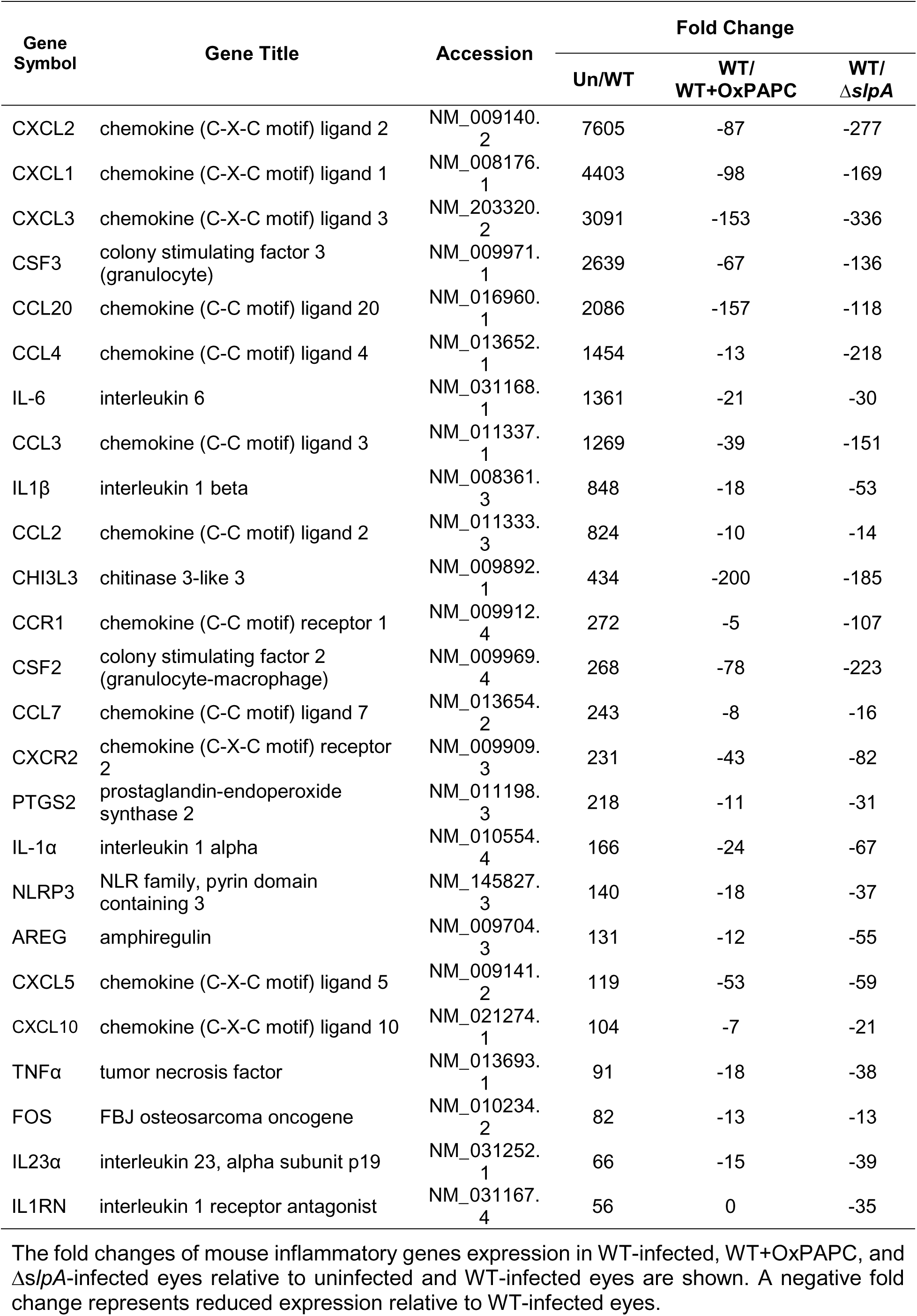
Top 25 differentially expressed mouse inflammatory genes in *Bacillus*-infected eyes at 10 hours postinfection.

Present treatment options, including intravitreal therapeutics and vitrectomy, are often unsuccessful at preventing irreversible damage to the retina that can result in total loss of vision. A complete understanding of the role of the host response in the disease process is required to design rational treatment strategies to improve the visual outcome of this disease. Infection with an SLP-deficient pathogen ^33^, chemical inhibition of TLR2 and TLR4 after infection ^34^, mice deficient in TLR2 ^36^, TLR4 ^37^, MyD88 ^38,85^, and TRIF^38^, mice pre-treated with TLR2 agonists ^77,99^, mice deficient in CXCL1^45^, and anti-CXCL1 administration ^45^ each resulted in reduced inflammation and better clinical outcomes. Collectively, these findings suggest targeting innate immune activation and their response elements as potential therapeutic strategies. Here, we identified several innate inflammatory genes which could be investigated further as potential anti-inflammatory targets in endophthalmitis. Further studies are necessary to determine whether expression of these genes contributes directly to the pathogenesis of *Bacillus* endophthalmitis. All things considered, this study demonstrated a viable strategy to minimize the otherwise robust and sight-threatening inflammation in *Bacillus* endophthalmitis and perhaps this infection caused by other pathogens.

## ACKNOWLEDGMENTS

The authors thank Dr. Agnes Fouet (Institut Cochin, INSERM U1016, CNRS UMR8104, University Paris Descartes, Paris, France) for providing the bacterial strains, Dr. Abdul Wadud Khan (Department of Surgical Oncology, The University of Texas MD Anderson Cancer Center) for generating the diversity index and heatmap, Dr. Feng Li and Mark Dittmar (OUHSC P30 Live Animal Imaging Core, Dean A. McGee Eye Institute, Oklahoma City, OK, USA), and the OUHSC P30 Cellular Imaging Core (Dean A. McGee Eye Institute, Oklahoma City, OK, USA) for histology expertise.

This work was supported by National Institutes of Health grants R01EY028810 and R01EY024140 (to MCC). Our research was also supported in part by National Institutes of Health grants R01EY025947 and R21EY028066 (to MCC), National Eye Institute Vision Core Grant P30EY021725 (to MCC), a Presbyterian Health Foundation Research Support Grant Award (to MCC and MHM), a Presbyterian Health Foundation Equipment Grant (to Robert E. Anderson, OUHSC), and an unrestricted grant to the Dean A. McGee Eye Institute from Research to Prevent Blindness. We thank the Laboratory for Molecular Biology and Cytometry Research at OUHSC for the use of the Core Facility which provided NanoString Sprint instrumentation, assay data, and analysis support.

## Notes

### Competing Interest Statement

The authors have declared no competing interest.

