## Supplemental Table 1 for "INNATE IMMUNE INTERFERENCE ATTENUATES INFLAMMATION IN *BACILLUS* ENDOPHTHALMITIS"

| **Supplementary Table 1. Differential expression of mouse inflammatory genes in Bacillus infected eyes at 10 hour postinfection** | | | | | |
| --- | --- | --- | --- | --- | --- |
| Gene Symbol | Gene title | Accession | Fold Change | | |
|  |  |  | Un/  WT | WT/  WT+OxPAPC | WT/  ∆*slpA* |
| CXCL2 | chemokine (C-X-C motif) ligand 2 | NM_009140.2 | 7605 | -87 | -277 |
| CXCL1 | chemokine (C-X-C motif) ligand 1 | NM_008176.1 | 4403 | -98 | -169 |
| CXCL3 | chemokine (C-X-C motif) ligand 3 | NM_203320.2 | 3091 | -153 | -336 |
| CSF3 | colony stimulating factor 3 (granulocyte) | NM_009971.1 | 2639 | -67 | -136 |
| CCL20 | chemokine (C-C motif) ligand 20 | NM_016960.1 | 2086 | -157 | -118 |
| CCL4 | chemokine (C-C motif) ligand 4 | NM_013652.1 | 1454 | -13 | -218 |
| IL-6 | interleukin 6 | NM_031168.1 | 1361 | -21 | -30 |
| CCL3 | chemokine (C-C motif) ligand 3 | NM_011337.1 | 1269 | -39 | -151 |
| IL-1β | interleukin 1 beta | NM_008361.3 | 848 | -18 | -53 |
| CCL2 | chemokine (C-C motif) ligand 2 | NM_011333.3 | 824 | -10 | -14 |
| CHI3L3 | chitinase 3-like 3 | NM_009892.1 | 434 | -200 | -185 |
| CCR1 | chemokine (C-C motif) receptor 1 | NM_009912.4 | 272 | -5 | -107 |
| CSF2 | colony stimulating factor 2 (granulocyte-macrophage) | NM_009969.4 | 268 | -78 | -223 |
| CCL7 | chemokine (C-C motif) ligand 7 | NM_013654.2 | 243 | -8 | -16 |
| CXCR2 | chemokine (C-X-C motif) receptor 2 | NM_009909.3 | 231 | -43 | -82 |
| PTGS2 | prostaglandin-endoperoxide synthase 2 | NM_011198.3 | 218 | -11 | -31 |
| IL1α | interleukin 1 alpha | NM_010554.4 | 166 | -24 | -67 |
| NLRP3 | NLR family, pyrin domain containing 3 | NM_145827.3 | 140 | -18 | -37 |
| AREG | amphiregulin | NM_009704.3 | 131 | -12 | -55 |
| CXCL5 | chemokine (C-X-C motif) ligand 5 | NM_009141.2 | 119 | -53 | -59 |
| CXCL10 | chemokine (C-X-C motif) ligand 10 | NM_021274.1 | 104 | -7 | -21 |
| TNFα | tumor necrosis factor | NM_013693.1 | 91 | -18 | -38 |
| FOS | FBJ osteosarcoma oncogene | NM_010234.2 | 82 | -13 | -13 |
| IL-23α | interleukin 23, alpha subunit p19 | NM_031252.1 | 66 | -15 | -39 |
| CEBPβ | CCAAT/enhancer binding protein (C/EBP), beta | NM_009883.3 | 55 | -5 | -16 |
| MAFF | v-maf musculoaponeurotic fibrosarcoma oncogene family, protein F (avian) | NM_010755.3 | 43 | -16 | -8 |
| IL-10 | interleukin 10 | NM_010548.1 | 30 | -6 | -9 |
| CCL11 | chemokine (C-C motif) ligand 11 | NM_011330.3 | 30 | -10 | -22 |
| TLR2 | toll-like receptor 2 | NM_011905.2 | 27 | -9 | -13 |
| CCL19 | chemokine (C-C motif) ligand 19 | NM_011888.2 | 22 | -6 | -6 |
| TNFSF14 | tumor necrosis factor (ligand) superfamily, member 14 | NM_019418.2 | 22 | -11 | -16 |
| CFB | complement factor B | NM_008198.2 | 20 | -10 | -14 |
| NOD2 | nucleotide-binding oligomerization domain containing 2 | NM_145857.2 | 18 | -7 | -7 |
| IIGP1 | interferon inducible GTPase 1 | NM_021792.3 | 12 | -2 | -9 |
| RELB | avian reticuloendotheliosis viral (v-rel) oncogene related B | NM_009046.2 | 12 | -2 | -5 |
| C3 | complement component 3 | NM_009778.2 | 10 | -7 | -11 |
| CXCR4 | chemokine (C-X-C motif) receptor 4 | NM_009911.3 | 8 | -2 | -7 |
| IFIT1 | interferon-induced protein with tetratricopeptide repeats 1 | NM_008331.2 | 7 | -2 | -5 |
| NF-kB1 | nuclear factor of kappa light polypeptide gene enhancer in B cells 1, p105 | NM_008689.2 | 7 | -4 | -5 |
| MYD88 | myeloid differentiation primary response gene 88 | NM_010851.2 | 6 | -2 | -4 |
| OAS2 | 2'-5' oligoadenylate synthetase 2 | NM_145227.2 | 6 | -3 | -5 |
| TYROBP | TYRO protein tyrosine kinase binding protein | NM_011662.2 | 5 | -3 | -5 |
| TLR4 | toll-like receptor 4 | NM_021297.2 | 5 | -3 | -6 |
| TNFαIP3 | tumor necrosis factor, alpha-induced protein 3 | NM_009397.2 | 4 | -3 | -4 |
| STAT3 | signal transducer and activator of transcription 3 | NM_213659.2 | 4 | -4 | -3 |
| IRF1 | interferon regulatory factor 1 | NM_008390.1 | 4 | -3 | -3 |
| HSPB1 | heat shock protein 1 | NM_013560.2 | 3 | -1 | -2 |
| ALOX12 | arachidonate 12-lipoxygenase | NM_007440.4 | 3 | -1 | -3 |
| IFI44 | interferon-induced protein 44 | NM_133871.2 | 2 | -2 | -2 |
| IL-10Rβ | interleukin 10 receptor, beta | NM_008349.5 | 2 | -2 | -3 |
| IFIT3 | interferon-induced protein with tetratricopeptide repeats 3 | NM_010501.1 | 2 | -1 | -2 |
| ARG1 | arginase, liver | NM_007482.3 | 2 | -1 | -2 |
| CFL1 | cofilin 1, non-muscle | NM_007687.5 | 2 | -1 | -1 |
| BIRC2 | baculoviral IAP repeat-containing 2 | NM_007465.2 | 1 | -1 | -1 |
| BCL2L1 | BCL2-like 1 | NM_009743.4 | 1 | -1 | -1 |
| CDC42 | cell division cycle 42 | NM_009861.1 | 1 | -1 | -1 |
| C1RA | complement component 1, r subcomponent A | NM_023143.3 | 1 | -1 | -1 |
| IRF7 | interferon regulatory factor 7 | NM_016850.2 | 1 | -1 | -1 |
| HIF1α | hypoxia inducible factor 1, alpha subunit | NM_010431.2 | 1 | -1 | -1 |
| MYL2 | myosin, light polypeptide 2, regulatory, cardiac, slow | NM_010861.3 | 1 | -1 | -1 |
| IFIT2 | interferon-induced protein with tetratricopeptide repeats 2 | NM_008332.2 | 1 | 0 | -1 |
| PRKCα | protein kinase C, alpha | NM_011101.3 | -1 | 1 | 1 |
| The fold changes of mouse inflammatory genes expression in WT-infected, WT+OxPAPC, and ∆*slpA*-infected eyes relative to uninfected and WT-infected eyes are shown. A negative fold change represents reduced expression relative to untreated WT-infected eyes. | | | | | |
