## Supplemental Table 2 for "INNATE IMMUNE INTERFERENCE ATTENUATES INFLAMMATION IN *BACILLUS* ENDOPHTHALMITIS"

| **Supplementary Table 2. Differential expression of mouse complement genes in Bacillus infected eyes at 10 hour postinfection** | | | | | |
| --- | --- | --- | --- | --- | --- |
| Gene symbol | Gene title | Accession | Fold Change | | |
|  |  |  | Un/  WT | WT/  WT+OxPAPC | WT/  ∆*slpA* |
| Cfb | complement factor B | NM_008198.2 | 19.6 | -10.1 | -13.5 |
| C3 | complement component 3 | NM_009778.2 | 9.9 | -6.8 | -10.8 |
| C6 | complement component 6 | NM_016704.2 | 2.9 | -1.2 | -3.3 |
| C7 | complement component 7 | XM_356827.6 | 2.6 | -0.4 | -2.0 |
| Hc | hemolytic complement | NM_010406.1 | 1.3 | -1.2 | -1.4 |
| C8α | complement component 8, alpha polypeptide | NM_146148.1 | 1.1 | -1.0 | -2.9 |
| C8b | complement component 8, beta polypeptide | NM_133882.2 | 1.1 | -0.6 | -0.3 |
| C1ra | complement component 1, r subcomponent A | NM_023143.3 | 1.0 | -0.5 | -0.9 |
| C1qb | complement component 1, q subcomponent, beta polypeptide | NM_009777.2 | 0.9 | -0.2 | -0.6 |
| C1s | complement component 1, s subcomponent | NM_144938.2 | 0.8 | -0.4 | -0.8 |
| C4a | complement component 4A (Rodgers blood group) | NM_011413.2 | 0.4 | -0.2 | -0.5 |
| C1qa | complement component 1, q subcomponent, alpha polypeptide | NM_007572.2 | 0.3 | -0.1 | -0.4 |
| C2 | complement component 2 (within H-2S) | NM_013484.2 | 0.1 | 0.0 | -0.1 |
| C3ar1 | complement component 3a receptor 1 | NM_009779.2 | -0.1 | -0.6 | -0.8 |
| C9 | complement component 9 | NM_013485.1 | -0.2 | -0.2 | 0.5 |
| Cfd | complement factor D (adipsin) | NM_013459.1 | -0.7 | -0.3 | -1.4 |
| The fold changes of mouse inflammatory genes expression in WT-infected, WT+OxPAPC, and ∆*slpA*-infected eyes relative to uninfected and WT-infected eyes are shown. A negative fold change represents reduced expression relative to untreated WT-infected eyes. | | | | | |
