## Supplemental Table 3 for "INNATE IMMUNE INTERFERENCE ATTENUATES INFLAMMATION IN *BACILLUS* ENDOPHTHALMITIS"

| **Supplementary Table 3. Differential expression of innate immune genes in *Bacillus* infected eyes at 10 hour postinfection** | | | | | |
| --- | --- | --- | --- | --- | --- |
| **Gene symbol** | **Gene Name** | **Accession** | **Fold Change** | | |
|  |  |  | **Un/**  **WT** | **WT/**  **WT+OxPAPC** | **WT/**  **∆*slpA*** |
| PTGS2 | prostaglandin-endoperoxide synthase 2 | NM_011198.3 | 218.0 | -10.8 | -31.1 |
| NLRP3 | NLR family, pyrin domain containing 3 | NM_145827.3 | 140.3 | -18.0 | -37.2 |
| TLR2 | toll-like receptor 2 | NM_011905.2 | 26.8 | -9.3 | -13.0 |
| TLR6 | toll-like receptor 6 | NM_011604.3 | 19.8 | -1.0 | -22.4 |
| NOD2 | nucleotide-binding oligomerization domain containing 2 | NM_145857.2 | 17.5 | -7.0 | -7.1 |
| RELB | avian reticuloendotheliosis viral (v-rel) oncogene related B | NM_009046.2 | 11.8 | -2.2 | -4.8 |
| TSLP | thymic stromal lymphopoietin | NM_021367.1 | 11.5 | -4.1 | -11.7 |
| LTB4R1 | leukotriene B4 receptor 1 | NM_008519.2 | 10.7 | -1.7 | -7.1 |
| NF-κB1 | nuclear factor of kappa light polypeptide gene enhancer in B cells 1, p105 | NM_008689.2 | 6.9 | -3.8 | -5.1 |
| MYD88 | myeloid differentiation primary response gene 88 | NM_010851.2 | 6.1 | -1.6 | -3.5 |
| NOS2 | nitric oxide synthase 2, inducible | NM_010927.3 | 5.9 | -2.1 | -5.8 |
| TLR8 | toll-like receptor 8 | NM_133212.2 | 5.5 | -0.3 | -3.3 |
| TYROBP | TYRO protein tyrosine kinase binding protein | NM_011662.2 | 5.3 | -3.4 | -5.3 |
| TLR4 | toll-like receptor 4 | NM_021297.2 | 5.0 | -2.7 | -5.7 |
| PTGIR | prostaglandin I receptor (IP) | NM_008967.3 | 4.8 | -1.2 | -8.2 |
| PTGS1 | prostaglandin-endoperoxide synthase 1 | NM_008969.3 | 4.4 | -1.4 | -4.4 |
| STAT3 | signal transducer and activator of transcription 3 | NM_213659.2 | 4.3 | -3.8 | -2.6 |
| PTGER4 | prostaglandin E receptor 4 (subtype EP4) | NM_008965.1 | 4.0 | -0.1 | -1.2 |
| RELA | v-rel reticuloendotheliosis viral oncogene homolog A (avian) | NM_009045.4 | 3.1 | -0.5 | -1.8 |
| HSPB1 | heat shock protein 1 | NM_013560.2 | 2.9 | -1.0 | -2.0 |
| LTB4R2 | leukotriene B4 receptor 2 | NM_020490.2 | 2.7 | -1.1 | -3.5 |
| TLR7 | toll-like receptor 7 | NM_133211.3 | 2.1 | -0.5 | -1.0 |
| TLR1 | toll-like receptor 1 | NM_030682.1 | 1.9 | -0.5 | -4.3 |
| MBL2 | mannose-binding lectin (protein C) 2 | NM_010776.1 | 1.0 | -0.8 | -1.4 |
| PTGER1 | prostaglandin E receptor 1 (subtype EP1) | NM_013641.2 | 1.0 | -0.3 | -0.8 |
| STAT2 | signal transducer and activator of transcription 2 | NM_019963.1 | 0.8 | -0.7 | -1.2 |
| STAT1 | signal transducer and activator of transcription 1 | NM_009283.3 | 0.6 | -0.4 | -0.8 |
| NOD1 | nucleotide-binding oligomerization domain containing 1 | NM_172729.2 | 0.4 | -0.5 | -0.8 |
| TLR3 | toll-like receptor 3 | NM_126166.2 | 0.4 | -0.2 | -0.3 |
| TLR9 | toll-like receptor 9 | NM_031178.2 | 0.4 | -1.0 | -0.5 |
| TOLLIP | toll interacting protein | NM_023764.3 | 0.4 | 0.0 | -0.2 |
| PDGFA | platelet derived growth factor, alpha | NM_008808.3 | 0.3 | -0.2 | -0.3 |
| TLR5 | toll-like receptor 5 | NM_016928.2 | 0.2 | 0.0 | -0.2 |
| HMGN1 | high mobility group nucleosomal binding domain 1 | NM_008251.3 | 0.1 | 0.0 | 0.0 |
| TWIST2 | twist basic helix-loop-helix transcription factor 2 | NM_007855.2 | 0.1 | 0.2 | 0.1 |
| HMGB2 | high mobility group box 2 | NM_008252.3 | 0.1 | 0.8 | 0.6 |
| HSPB2 | heat shock protein 2 | NM_024441.3 | 0.0 | 0.0 | 0.0 |
| PTGER2 | prostaglandin E receptor 2 (subtype EP2) | NM_008964.4 | 0.0 | -0.3 | -1.3 |
| HMGB1 | high mobility group box 1 | NM_010439.3 | -0.1 | 0.1 | 0.1 |
| PTGFR | prostaglandin F receptor | NM_008966.3 | -0.2 | -0.1 | 0.1 |
| PTGER3 | prostaglandin E receptor 3 (subtype EP3) | NM_011196.2 | -0.3 | 0.1 | -0.3 |
| The fold changes of mouse inflammatory genes expression in WT-infected, WT+OxPAPC, and ∆s*lpA*-infected eyes relative to uninfected and WT-infected eyes are shown. A negative fold change represents reduced expression relative to untreated WT-infected eyes. | | | | | |
