## Supplemental Table 4 for "INNATE IMMUNE INTERFERENCE ATTENUATES INFLAMMATION IN *BACILLUS* ENDOPHTHALMITIS"

| **Supplementary Table 4. Differential expression of inflammatory cytokines in *Bacillus* infected eyes at 10 hour postinfection** | | | | | |
| --- | --- | --- | --- | --- | --- |
| **Gene symbol** | **Gene title** | **Accession** | **Fold Change** | | |
|  |  |  | **Un/**  **WT** | **WT/**  **WT+OxPAPC** | **WT/**  **∆*slpA*** |
| CSF3 | colony stimulating factor 3 (granulocyte) | NM_009971.1 | 2639.2 | -67.5 | -136.3 |
| IL-6 | interleukin 6 | NM_031168.1 | 1360.9 | -21.1 | -29.7 |
| IL-1β | interleukin 1 beta | NM_008361.3 | 848.3 | -18.2 | -53.1 |
| CSF2 | colony stimulating factor 2 (granulocyte-macrophage) | NM_009969.4 | 268.0 | -77.8 | -222.8 |
| IL-1α | interleukin 1 alpha | NM_010554.4 | 166.4 | -24.0 | -66.7 |
| TNFα | tumor necrosis factor | NM_013693.1 | 90.9 | -18.0 | -38.2 |
| IL-23α | interleukin 23, alpha subunit p19 | NM_031252.1 | 66.0 | -15.1 | -39.1 |
| IL1RN | interleukin 1 receptor antagonist | NM_031167.4 | 56.3 | -0.4 | -35.4 |
| IL18RAP | interleukin 18 receptor accessory protein | NM_010553.2 | 35.2 | -0.5 | -35.2 |
| IL-10 | interleukin 10 | NM_010548.1 | 30.2 | -5.5 | -8.9 |
| TNFSF14 | tumor necrosis factor (ligand) superfamily, member 14 | NM_019418.2 | 21.8 | -10.8 | -15.5 |
| IIGP1 | interferon inducible GTPase 1 | NM_021792.3 | 12.1 | -2.3 | -9.4 |
| IFIT1 | interferon-induced protein with tetratricopeptide repeats 1 | NM_008331.2 | 7.2 | -2.1 | -5.4 |
| IL-11 | interleukin 11 | NM_008350.2 | 6.4 | -1.3 | -4.3 |
| TNFαip3 | tumor necrosis factor, alpha-induced protein 3 | NM_009397.2 | 4.5 | -3.0 | -4.0 |
| IRF1 | interferon regulatory factor 1 | NM_008390.1 | 3.5 | -3.1 | -3.2 |
| IL-6Rα | interleukin 6 receptor, alpha | NM_010559.2 | 3.4 | -0.5 | -1.8 |
| IL-23R | interleukin 23 receptor | NM_144548.1 | 3.3 | -0.8 | -2.5 |
| TGFβ 1 | transforming growth factor, beta 1 | NM_011577.1 | 3.2 | -1.0 | -2.9 |
| IRF5 | interferon regulatory factor 5 | NM_012057.3 | 2.9 | -1.2 | -2.6 |
| IL-5 | interleukin 5 | NM_010558.1 | 2.5 | 0.1 | -1.3 |
| IL-12β | interleukin 12b | NM_008352.1 | 2.4 | -0.7 | -1.2 |
| IFI44 | interferon-induced protein 44 | NM_133871.2 | 2.3 | -1.9 | -2.0 |
| IL-17A | interleukin 17A | NM_010552.3 | 2.3 | -0.5 | -1.3 |
| IL-10Rβ | interleukin 10 receptor, beta | NM_008349.5 | 2.3 | -1.6 | -2.9 |
| IL-22Rα2 | interleukin 22 receptor, alpha 2 | NM_178258.5 | 2.3 | 0.1 | -2.7 |
| IL-1R1 | interleukin 1 receptor, type I | NM_001123382.1 | 2.1 | -0.2 | -1.3 |
| IFNB1 | interferon beta 1, fibroblast | NM_010510.1 | 2.1 | -0.3 | -0.5 |
| IFIT3 | interferon-induced protein with tetratricopeptide repeats 3 | NM_010501.1 | 1.8 | -1.1 | -2.0 |
| IL-1RAP | interleukin 1 receptor accessory protein | NM_008364.2 | 1.8 | -0.3 | -2.3 |
| IL-12α | interleukin 12a | NM_008351.1 | 1.8 | -0.8 | -2.5 |
| IL-3 | interleukin 3 | NM_010556.4 | 1.7 | -0.9 | -1.5 |
| I-L2 | interleukin 2 | NM_008366.3 | 1.5 | -0.1 | -1.1 |
| IL-13 | interleukin 13 | NM_008355.2 | 1.4 | -1.5 | -0.4 |
| TGFβR1 | transforming growth factor, beta receptor I | NM_009370.2 | 1.4 | -0.2 | -0.9 |
| IRF7 | interferon regulatory factor 7 | NM_016850.2 | 1.0 | -1.0 | -0.7 |
| IL-22 | interleukin 22 | NM_016971.1 | 0.9 | 0.0 | -0.3 |
| IRF3 | interferon regulatory factor 3 | NM_016849.3 | 0.9 | -0.3 | -1.1 |
| TRAF2 | TNF receptor-associated factor 2 | NM_009422.2 | 0.8 | -0.5 | -1.2 |
| IFN𝛾 | interferon gamma | NM_008337.1 | 0.7 | 0.4 | 0.4 |
| IFIT2 | interferon-induced protein with tetratricopeptide repeats 2 | NM_008332.2 | 0.7 | -0.5 | -0.8 |
| IL-7 | interleukin 7 | NM_008371.2 | 0.6 | 0.0 | -1.3 |
| IL15 | interleukin 15 | NM_008357.1 | 0.5 | -1.5 | -0.8 |
| IL-21 | interleukin 21 | NM_021782.2 | 0.3 | -0.1 | -0.1 |
| IL-4 | interleukin 4 | NM_021283.1 | 0.1 | 0.3 | -0.2 |
| TGFβ2 | transforming growth factor, beta 2 | NM_009367.1 | 0.1 | -0.1 | -0.1 |
| IL-18 | interleukin 18 | NM_008360.1 | 0.1 | -0.1 | -0.1 |
| IFNA1 | interferon alpha 1 | NM_010502.2 | -0.1 | 0.4 | 0.4 |
| IL-9 | interleukin 9 | NM_008373.1 | -0.2 | 0.2 | -0.6 |
| IFI27L2A | interferon, alpha-inducible protein 27 like 2A | NM_029803.1 | -0.4 | -0.1 | -0.4 |
| TGFβ3 | transforming growth factor, beta 3 | NM_009368.2 | -0.5 | 0.2 | 0.4 |
| The fold changes of mouse inflammatory genes expression in WT-infected, WT+OxPAPC, and ∆s*lpA*-infected eyes relative to uninfected and WT-infected eyes are shown. A negative fold change represents reduced expression relative to untreated WT-infected eyes. | | | | | |
