## Supplemental Table 5 for "INNATE IMMUNE INTERFERENCE ATTENUATES INFLAMMATION IN *BACILLUS* ENDOPHTHALMITIS"

| **Supplementary Table 5. Differential expression of chemokines in *Bacillus* infected eyes at 10 hour postinfection** | | | | | |
| --- | --- | --- | --- | --- | --- |
| **Gene symbol** | **Gene title** | **Accession** | **Fold Change** | | |
|  |  |  | **Un/**  **WT** | **WT/**  **WT+OxPAPC** | **WT/**  **∆*slpA*** |
| CXCL2 | chemokine (C-X-C motif) ligand 2 | NM_009140.2 | 7605 | -87 | -277 |
| CXCL1 | chemokine (C-X-C motif) ligand 1 | NM_008176.1 | 4403 | -98 | -169 |
| CXCL3 | chemokine (C-X-C motif) ligand 3 | NM_203320.2 | 3091 | -153 | -336 |
| CCL20 | chemokine (C-C motif) ligand 20 | NM_016960.1 | 2086 | -157 | -118 |
| CCL4 | chemokine (C-C motif) ligand 4 | NM_013652.1 | 1454 | -13 | -218 |
| CCL3 | chemokine (C-C motif) ligand 3 | NM_011337.1 | 1269 | -39 | -151 |
| CCL2 | chemokine (C-C motif) ligand 2 | NM_011333.3 | 824 | -10 | -14 |
| CCR1 | chemokine (C-C motif) receptor 1 | NM_009912.4 | 272 | -5 | -107 |
| CCL7 | chemokine (C-C motif) ligand 7 | NM_013654.2 | 243 | -8 | -16 |
| CXCR2 | chemokine (C-X-C motif) receptor 2 | NM_009909.3 | 231 | -43 | -82 |
| CXCL5 | chemokine (C-X-C motif) ligand 5 | NM_009141.2 | 119 | -53 | -59 |
| CXCL10 | chemokine (C-X-C motif) ligand 10 | NM_021274.1 | 104 | -7 | -21 |
| CCL11 | chemokine (C-C motif) ligand 11 | NM_011330.3 | 30 | -10 | -22 |
| CXCL9 | chemokine (C-X-C motif) ligand 9 | NM_008599.2 | 28 | -2 | -14 |
| CCL19 | chemokine (C-C motif) ligand 19 | NM_011888.2 | 22 | -6 | -6 |
| CCL17 | chemokine (C-C motif) ligand 17 | NM_011332.2 | 17 | 0 | -3 |
| CCL5 | chemokine (C-C motif) ligand 5 | NM_013653.1 | 11 | -2 | -11 |
| CCR7 | chemokine (C-C motif) receptor 7 | NM_007719.2 | 11 | 0 | -5 |
| CXCR4 | chemokine (C-X-C motif) receptor 4 | NM_009911.3 | 8 | -2 | -7 |
| CCL24 | chemokine (C-C motif) ligand 24 | NM_019577.4 | 6 | -1 | -1 |
| CXCR1 | chemokine (C-X-C motif) receptor 1 | NM_178241.4 | 3 | 0 | -1 |
| CCR2 | chemokine (C-C motif) receptor 2 | NM_009915.2 | 2 | -5 | -7 |
| CCR3 | chemokine (C-C motif) receptor 3 | NM_009914.4 | 2 | 0 | -2 |
| CCL22 | chemokine (C-C motif) ligand 22 | NM_009137.2 | 1 | 0 | 0 |
| CCL8 | chemokine (C-C motif) ligand 8 | NM_021443.2 | 1 | 0 | -2 |
| CCL21A | chemokine (C-C motif) ligand 21A (serine) | NM_011124.4 | 0 | 0 | 0 |
| CCR4 | chemokine (C-C motif) receptor 4 | NM_009916.2 | 0 | 1 | 1 |
| The fold changes of mouse inflammatory genes expression in WT-infected, WT+OxPAPC, and ∆s*lpA*-infected eyes relative to uninfected and WT-infected eyes are shown. A negative fold change represents reduced expression relative to untreated WT-infected eyes. | | | | | |
